# A class act: HDAC1-*Malat1* regulates MDSC apoptosis and cell cycling to decrease suppression of T cells

**DOI:** 10.64898/2026.03.23.713743

**Authors:** Aaron G. Baugh, Yingtong Liu, Edgar Gonzalez, Batul Al-Zubeidy, Mukund Iyer, Alexander H. Lee, Nana Gyabaah-Kessie, Matthew B. Jacobo, Cheol Park, Jesse Kreger, Lori Khatchaturian, Sabrina K. Zhong, Kaliya Acevedo, Saul J. Priceman, Josh Neman, Adam L. MacLean, Evanthia T. Roussos Torres

## Abstract

Myeloid derived suppressor cells (MDSCs) are key players in the immune-suppressed tumor microenvironment (TME) and significantly contribute to immune checkpoint inhibition (ICI) resistance, making them favorable targets for cancer immunotherapy. Epigenetic reprogramming of MDSCs using histone deacetylase (HDAC) inhibitors shows promise to sensitize the TME to ICIs. However, the molecular mechanism of HDAC inhibition in MDSCs has yet to be elucidated. Murine and human MDSC models treated with Entinostat revealed that the long non-coding RNA *Malat1* downregulates pSTAT3 and decreases MDSC-mediated suppression of T cell proliferation. Through HDAC inhibitor screens, we identified HDAC1 as preferentially regulating *Malat1* expression, STAT3 activation, and MDSC suppression. We also show that HDAC1 inhibition increases MDSC apoptosis by shifting pro-vs. anti-apoptotic signals and increases G0/G1 cell cycle arrest via decreasing G1-S transition cyclin-CDK complexes. Collectively, our findings provide a multi-pronged mechanism of HDAC inhibition in MDSCs that inform the development of future rational combination therapies.

**One Sentence Summary:** HDAC1 inhibition in MDSCs increases *Malat1*, decreases pSTAT3, induces apoptosis/cell cycle arrest, and decreases suppression of T cells

## INTRODUCTION

Myeloid derived suppressor cells (MDSCs) are a heterogenous population of immature neutrophils and monocytes with robust immunosuppressive activity to inhibit T cell, B cell, and natural killer (NK) cell immune responses (1). They are highly present in various disease settings with prolonged myeloid growth factors and inflammatory signals, such as in cancers, chronic infections, aging, and autoimmune diseases (2–5). In the context of cancer, MDSCs have been reported to support tumor growth and survival, promote angiogenesis, facilitate metastasis, and enable immune evasion, ultimately contributing to poor prognosis and exacerbation of advanced-stage disease (6,7). Solid tumors, such as breast cancer, are characterized by extensive MDSC infiltration in the tumor microenvironment (TME), both at primary and metastatic sites (8), which interfere with immune checkpoint inhibitor (ICI) therapies (9,10). Given the central role of MDSCs in creating a suppressive TME, efforts to modulate MDSC function and sensitize the TME to increase ICI-responsiveness through combination therapies are gaining traction.

Epigenetic agents that remodel chromatin states, such as histone deacetylase (HDAC) inhibitors, have been previously studied with ICIs to alter immune responses in the TME (11). Our group and others have shown that the combination of Entinostat, a class I HDAC inhibitor, with anti-PD-1 and anti-CTLA-4 increased survival and decreased metastases in murine models of breast cancer (11,12) and yielded a 25% objective response rate in patients with metastatic HER2-negative breast cancer from the Phase 1b trial NCI-9844 (13,14). While HDAC inhibitors have been more extensively studied in directly inhibiting cancer cells, an increasing area of interest is the modulatory effects on suppressive immune cell populations (15,16). Previous studies have shown that HDAC inhibitors can decrease MDSC suppressive function, expansion, and recruitment to tumor sites (12,17,18), but the specific molecular mechanisms of HDAC inhibition in MDSCs have yet to be fully elucidated.

In this study, through statistical modeling to infer biologically interpretable marker genes in MDSCs and integrated epigenome and transcriptome analyses, we discovered a coordinated mechanism through which HDAC inhibition leads to decreased MDSC function. We provide evidence in murine and human MDSC models that HDAC inhibition upregulates long non-coding RNA *Malat1*, decreases STAT3 signaling, increases apoptosis, and increases cell cycle arrest to decrease suppression of T cells. Our findings present multiple actionable targets for combination immunotherapies to overcome MDSC suppression and improve cancer treatment outcomes.

## RESULTS

### MDSC-mediated suppression of T cells is HDAC1-dependent

We have previously shown that pre-treatment of MDSC-like J774M cell line with Entinostat, a class I HDAC inhibitor preferentially targeting HDACs 1/3 with minimal inhibition of HDAC2, reduces their suppressive function to restore T cell proliferation (17). Here, we show that Entinostat decreases MDSC suppression when co-cultured with CFSE proliferation dye-stained murine CD8+ T cells at 5 uM, a 10-fold lower dose than previously used. (**Fig. 1A**). We observe similar findings of decreased suppression in human PBMC-derived MDSCs pre-treated with Entinostat and co-cultured with CFSE stained human CD3+ T cells (**Fig. S1 and Fig. 1B**). Even though Entinostat has shown clinical benefit in combination with anti-PD-1 and anti-CTLA-4 ICIs, we were interested in examining whether even more specific HDAC-targeting inhibitors would yield similar effects on MDSC suppression. Thus, we pre-treated J774M cells with Chlopynostat (HDAC1 inhibitor), Santacruzamate A (an HDAC2 inhibitor), RGFP966 (an HDAC3 inhibitor), and TMP269 (a class IIa HDAC inhibitor targeting HDACs 4, 5, 7, and 9), before co-culturing with CD8+ T cells. We found that of the class I HDACs, inhibition of HDAC1 with Chlopynostat most robustly decreased MDSC suppression of T cells, whereas inhibition of HDAC2, HDAC3, or class IIa HDACs provided little or no reversal of suppression (**Fig. 1C**). Our findings provide evidence suggesting that HDAC1 may be preferentially responsible for regulating T cell suppression by MDSCs.

**Figure 1:**
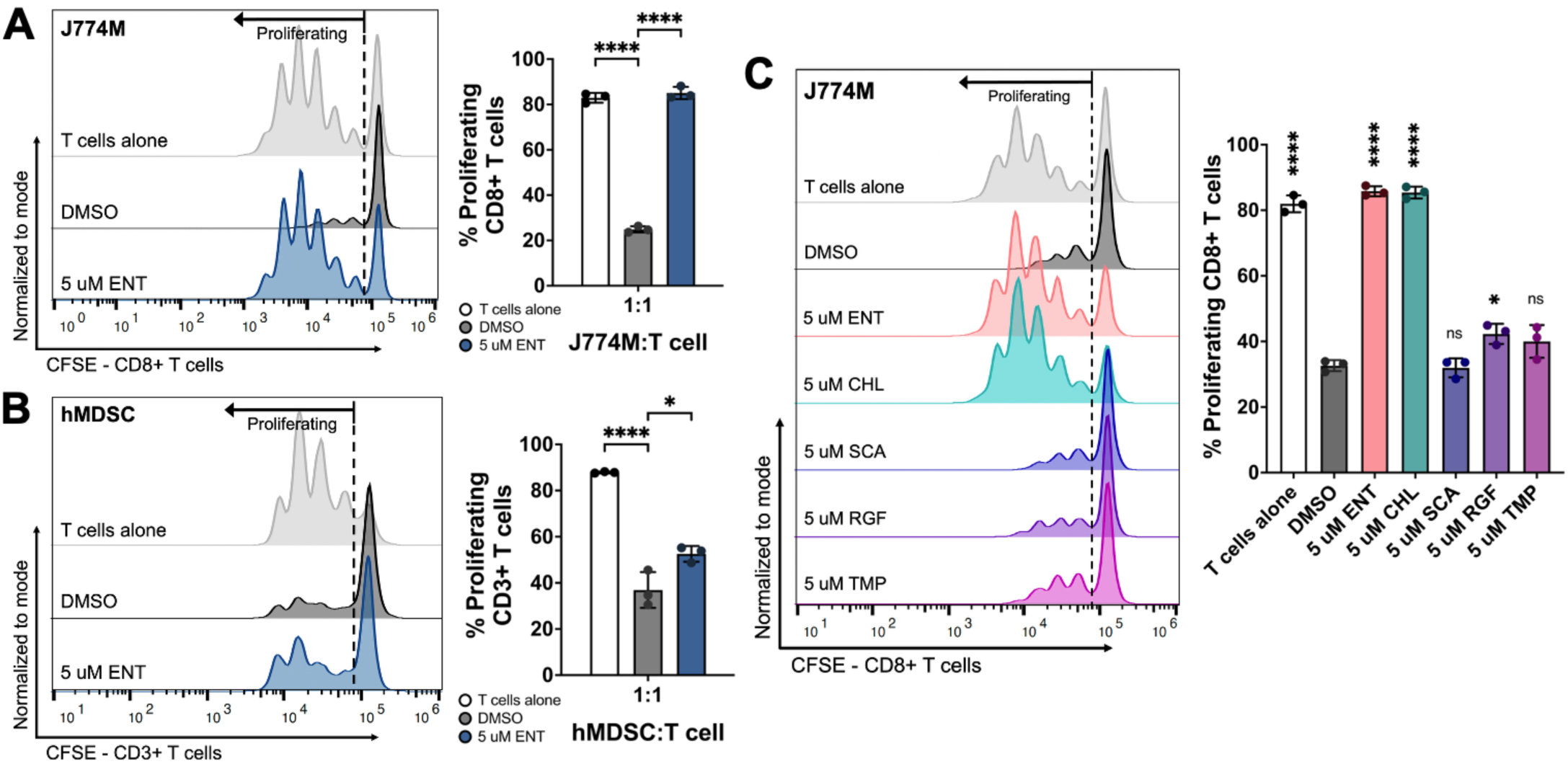
MDSC suppression of T cells is HDAC1-dependent. **(A)** Representative flow cytometry plot (left) and quantification (right) of proliferating murine CD8+ T cells after 66-72 hours of co-culture in a 1:1 ratio with J774M cells pre-treated for 72 hours with Entinostat (n=3). **(B)** Representative flow cytometry plot (left) and quantification (right) of proliferating human CD3+ T cells after 66-72 hours of co-culture in a 1:1 ratio with hMDSCs pre-treated for 48 hours with Entinostat (n=3). **(C)** Representative flow cytometry plot (left) and quantification (right) of proliferating murine CD8+ T cells after 66-72 hours of co-culture in a 1:1 ratio with J774M cells pre-treated for 72 hours with various HDAC inhibitors (n=3). One-way ANOVA, ns = not significant, * p<0.05, **** p<0.0001. ENT = Entinostat (HDAC1/2/3i); CHL = Chlopynostat (HDAC1i); SCA = Santacruzamate A (HDAC2i); RGF = RGPF966 (HDAC3i); TMP = TMP269 (class IIa HDACi, HDAC4/5/7/9i).

### HDAC1 inhibition increases lncRNA Malat1 expression in MDSCs

Our previous work in cancer systems immunology examined the complexities in cellular interactions and immune functions brought about by HDAC inhibition (Entinostat [E]) in combination with dual ICIs (anti-PD-1 [P] + anti-CTLA-4 [C]), referred to as EPC hereafter, in the breast-to-lung metastatic TME (19). We identified various cell states from single-cell RNA sequencing (scRNA-seq) of metastatic lung tumors in the NT2.5-LM (20) murine model of breast cancer, including immature myeloid-derived suppressor cells (MDSCs) and their two major cell states: granulocytic (G)- and monocytic (M)-MDSCs (**Fig. 2A**). Given the significant contribution of MDSCs to an immune-suppressed TME and previous work demonstrating decreased MDSC suppressive function upon treatment with Entinostat (12,17), we sought to uncover molecular mechanisms of HDAC inhibition in MDSCs. To identify a small, biologically informative gene set that captures subtle and complex transcriptional differences initiated by HDAC inhibition, we applied iterative logistic regression (iLR) (21) on the MDSC cluster and compared vehicle (V) with the EPC combination treatment group that yielded pre-clinical and clinical survival benefit in patients with metastatic breast cancer (12–14). iLR-identified genes marking for treatment with EPC were found to be involved in myeloid suppressive function, myeloid differentiation, metabolism, and ubiquitin-related pathways (22–39) (**Fig. 2B**). Of note, iLR identified *Malat1*, which was previously shown in MDSCs to target phosphorylated STAT3 (pSTAT3) for ubiquitin-mediated degradation (40). *S100a9* is a known direct target of pSTAT3 (41), and we previously showed that Entinostat decreased pSTAT3 expression in MDSCs (17). Together, these observations led us to ask whether HDAC inhibition in MDSCs acts upstream of STAT3 activation by upregulating *Malat1*.

**Figure 2:**
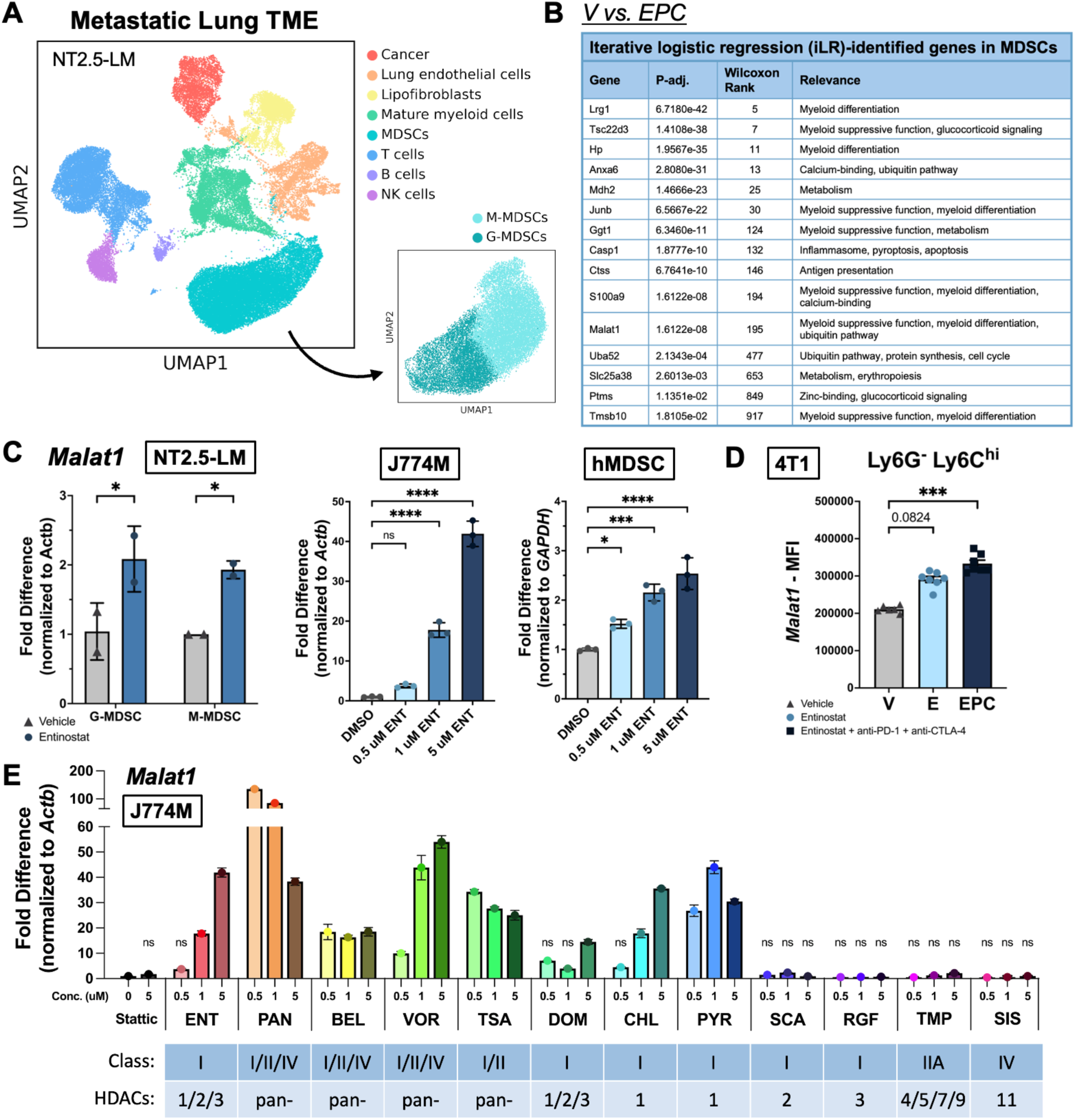
HDAC1 inhibition increases lncRNA Malat1 expression in MDSCs. **(A)** UMAP visualization of major cell clusters identified from single cell RNA sequencing (scRNA-seq) of NT2.5-LM breast-to-lung metastatic tumors after 3 weeks of treatment with: vehicle (V), Entinostat (E), anti-CTLA-4 + anti-PD-1 immune checkpoint inhibitors (PC), and Entinostat + anti-CTLA-4 + anti-PD-1 combination (EPC). Subclusters of MDSCs identified: G-MDSCs and M-MDSCs. **(B)** Iterative logistic regression analyses conducted on the major MDSC cluster comparing the Vehicle (V) and Entinostat + ICIs combination treatment (EPC). **(C)** Expression of *Malat1*/*MALAT1* in G- and M-MDSCs isolated from lung metastases of NT2.5-LM mice treated with vehicle vs. Entinostat for 3 weeks (left, n=2 from 5 pooled mice per treatment group), J774M murine MDSC-like cell line treated with Entinostat for 24 hours (middle, n=3), and human PBMC-derived MDSCs (hMDSCs) treated with Entinostat for 24 hours (right, n=3). **(D)** Median Fluorescence Intensity expression of *Malat1* in CD45^+^CD11b^+^ MHC-II^-^F4/80^-^Ly6G^-^Ly6C^hi^ cells from lung metastases in 4T1 mice, treated with vehicle (V), Entinostat (E), and Entinostat + ICIs (EPC) for 3 weeks. **(E)** Expression of *Malat1* in J774M cell line treated with various concentrations of various HDAC inhibitors, with corresponding targeted class and specific HDACs. One-way ANOVA for (C, E: all statistically significant with p<0.05, unless indicated as non-significant (ns)), Kruskal-Wallis test with Dunn’s correction for (D). * p<0.05, *** p< 0.001, **** p<0.0001. ENT = Entinostat; PAN = Panobinostat; BEL = Belinostat; VOR = Vorinostat; TSA = Trichostatin A; DOM = Domatinostat; CHL = Chlopynostat; PYR = Pyroxamide; SCA = Santacruzamate A; RGF = RGFP966; TMP = TMP269; SIS = SIS17.

We found that Entinostat treatment increased the expression of *Malat1* in G-MDSCs and M-MDSCs isolated from metastatic lung tumors in the NT2.5-LM model of breast cancer, whereby mice were treated with vehicle or Entinostat for 3 weeks (**Fig. 2C**). In both J774M cells and hMDSCs, we also found that Entinostat treatment increased the expression of *Malat1* and *MALAT1*, respectively (**Fig. 2C**). Given that four out of the five responding patients from the NCI-9844 trial had triple negative breast cancer (TNBC) (13), we also investigated the effects of combination treatment on *Malat1* expression in the 4T1 murine model of TNBC. After 3 weeks of Entinostat treatment in 4T1 lung metastases-bearing mice, we performed PrimeFlow cytometry staining to evaluate *Malat1* expression in MDSCs (CD45^+^CD11b^+^MHC-II^-^F4-80^-^). In the M-MDSC (Ly6G^-^Ly6C^hi^) population, we found that Entinostat treatment increased the median fluorescence intensity (MFI) expression of *Malat1* (**Fig. 2D**). Our findings in multiple systems suggest that HDAC inhibition significantly and dose-dependently increases *Malat1* expression in MDSCs.

We also found that HDAC inhibitors targeting HDAC1 specifically increased expression of *Malat1* in MDSCs, whereas those not targeting HDAC1 did not, by performing an HDAC inhibitor screen in J774M cells with various pan-(Panobinostat, Belinostat, Vorinostat, Trichostatin A), class I-(Entinostat, Domatinostat), HDAC1-(Chlopynostat, Pyroxamide), HDAC2-(Santacruzamate A), HDAC3-(RGFP966), class IIa-(TMP269), and class IV-(SIS17) HDAC inhibitors (**Fig. 2E**). Collectively, from unsupervised iterative logistic regression analyses and gene expression validations, we were able to infer a biologically relevant mechanism of HDAC1 inhibition in MDSCs upregulating *Malat1* to modulate immune pathway signaling.

### HDAC1 inhibition decreases STAT3 pathway activation

The STAT3 pathway has been shown to be involved in regulating MDSC expansion, survival, and function (42), and our previous work in the primary breast TME suggested that Entinostat decreases STAT3 activation (20). To determine if HDAC inhibition also decreases STAT3 activation in the lung metastatic TME, we established lung metastatic tumors in the NT2.5-LM murine model of breast cancer, and after 2 weeks of Entinostat treatment, isolated Ly6G^+^ G-MDSCs from collected lungs. We found that G-MDSCs from the lung metastases of Entinostat-treated mice had decreased pSTAT3 expression compared to those from vehicle-treated mice (**Fig. 3A**). Cell circuit analysis in the lung metastatic TME previously identified a decrease in interferon gamma (IFN-γ) signaling between G-MDSCs and CD8+ T cells upon HDAC inhibition (19), and we found that Entinostat treatment of J774M cells with IFN-γ stimulation also decreased pSTAT3 expression (**Fig. 3B**). In hMDSCs, Entinostat dose-dependently decreased pSTAT3 expression (**Fig. 3C**).

**Figure 3:**
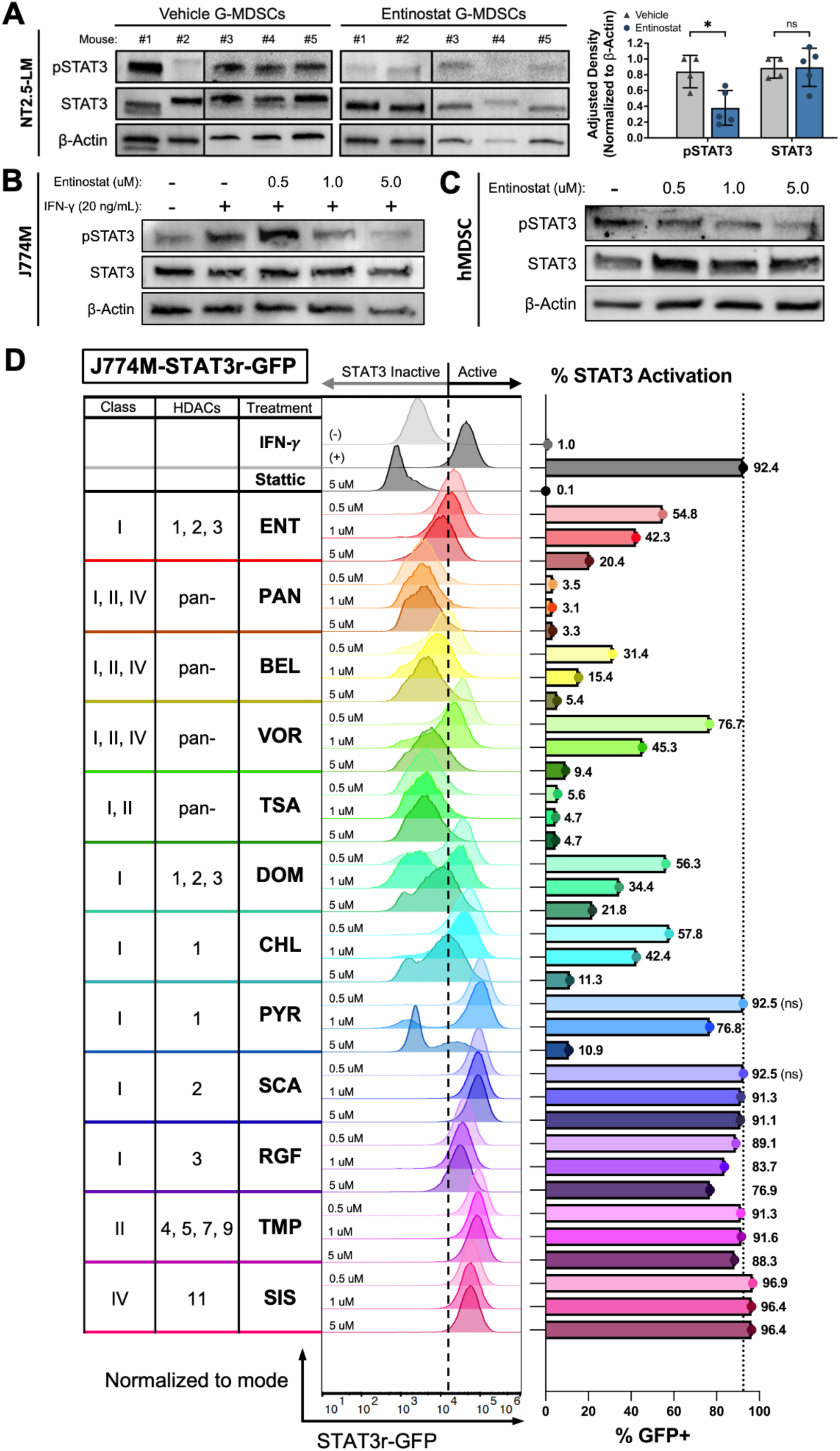
HDAC1 inhibition decreases STAT3 pathway activation in MDSCs. **(A)** Western blots of Tyr705 phosphorylated-STAT3 (pSTAT3), STAT3, and Beta-actin in G-MDSCs isolated from NT2.5-LM metastatic lung tumors. G-MDSCs were isolated with Ly-6G+ magnetic beads from single cell suspensions of dissociated metastatic lung tumors and stimulated with LPS (2 hours, 1 ug/mL). NT2.5-LM tumor-bearing mice were treated with vehicle vs. Entinostat (5 mg/kg, 5x/wk) for 3 weeks. Each lane represents one mouse (n=5 per treatment group). 50 ug of total protein lysate loaded in each lane. Band density quantified with Image J and adjusted density normalized to Beta-actin (right), outlier mouse #2 in Vehicle removed with ROUT at threshold Q=1%. Two-way ANOVA, * p<0.05. **(B)** Western blot of pSTAT3, STAT3, and Beta-actin in J774M cells after 24 hours of Entinostat treatment and IFN-γ stimulation (30 min, 20 ng/mL). 20 ug of total protein lysate loaded in each lane. **(C)** Western blot of pSTAT3, STAT3, and Beta-actin in hMDSCs after 24 hours of Entinostat treatment. 50 ug of total protein lysate loaded in each lane. **(D)** Representative flow plots (left) and quantification (right, n=3) of J774M STAT3-responsive GFP reporter after 18 hours of pre-treatment with various concentrations of various HDAC inhibitors and IFN-γ stimulation for 5-7 hours (20 ng/mL), indicative of percentage of STAT3 activation. One-way ANOVA for (D: all statistically significant with p<0.05, unless indicated as non-significant (ns)). ENT = Entinostat; PAN = Panobinostat; BEL = Belinostat; VOR = Vorinostat; TSA = Trichostatin A; DOM = Domatinostat; CHL = Chlopynostat; PYR = Pyroxamide; SCA = Santacruzamate A; RGF = RGFP966; TMP = TMP269; SIS = SIS17.

To determine if the observed decrease in STAT3 activation is also HDAC1-dependent, we generated a STAT3-responsive GFP-reporter J774M cell line, in which 4 repeats of the M67 STAT3-specific DNA binding sequence was inserted in front of the GFP coding sequence (**Fig. S2**). Upon immune stimulation and STAT3 activation, GFP expression is induced. Using this reporter, we performed the same HDAC inhibitor screen and found that HDAC1 inhibitors most robustly decreased %GFP+ cells, indicative of decreased STAT3 activation (**Fig. 3D**). In sum, we show that HDAC inhibition, and specifically via HDAC1, dose-dependently decreases activated STAT3 in our murine and human MDSC models.

### HDAC1/3 inhibition decreases anti-apoptotic, cell cycle progression, and TNF signaling

To gain an understanding into signaling pathways altered by HDAC inhibition, we performed bulk RNA sequencing (bulkRNA-seq) on IFN-γ stimulated J774M cells treated with vehicle vs. Entinostat. Differential gene expression analysis by DESeq2 identified 2055 up-regulated and 1940 down-regulated genes upon Entinostat treatment (p-adj. < 0.05 and |log_2_FC| > 1.0; **Table S1**). Gene set enrichment analyses (GSEA) of differentially expressed genes revealed that Entinostat treatment significantly downregulated “Cytokine-cytokine receptor interaction”, “JAK-STAT signaling”, “NF-kappa B signaling”, “Apoptosis”, “DNA replication”, “Cell Cycling”, and “TNF signaling” pathways (**Fig. 4A, S3A-F**). We confirmed an increase in *Malat1* expression (7.18-fold, p-adj. = 3.44E-07), and found significant decreases in the expression of TNF/NF-κB signaling genes (*Il6, Tnf, Fas, Nfkb1, Nfkb2, Relb, Myd88, Tnfsf10/*TRAIL), anti-apoptotic Bcl-2 family genes (*Bcl2, Bcl2a1a, Bcl2a1b, Bcl2a1d, Bcl2l1*), microtubule genes governing apoptosis (*Tuba1b, Tuba1c, Tuba4a*), cyclins (*Ccna2, Ccnb1, Ccnb2, Ccnd1, Ccne1*) and cyclin dependent kinases (*Cdk1, Cdk2, Cdk4, Cdk6*), and various cell cycle regulatory genes (*Mcm2-6*, *Cdc6, Cdc20, Cdc25a, Cdc45, E2f1, Chek1*). Since HDAC inhibition redistributes histone acetylation patterns to affect transcription, we were interested in examining differentially active promoters.

**Figure 4:**
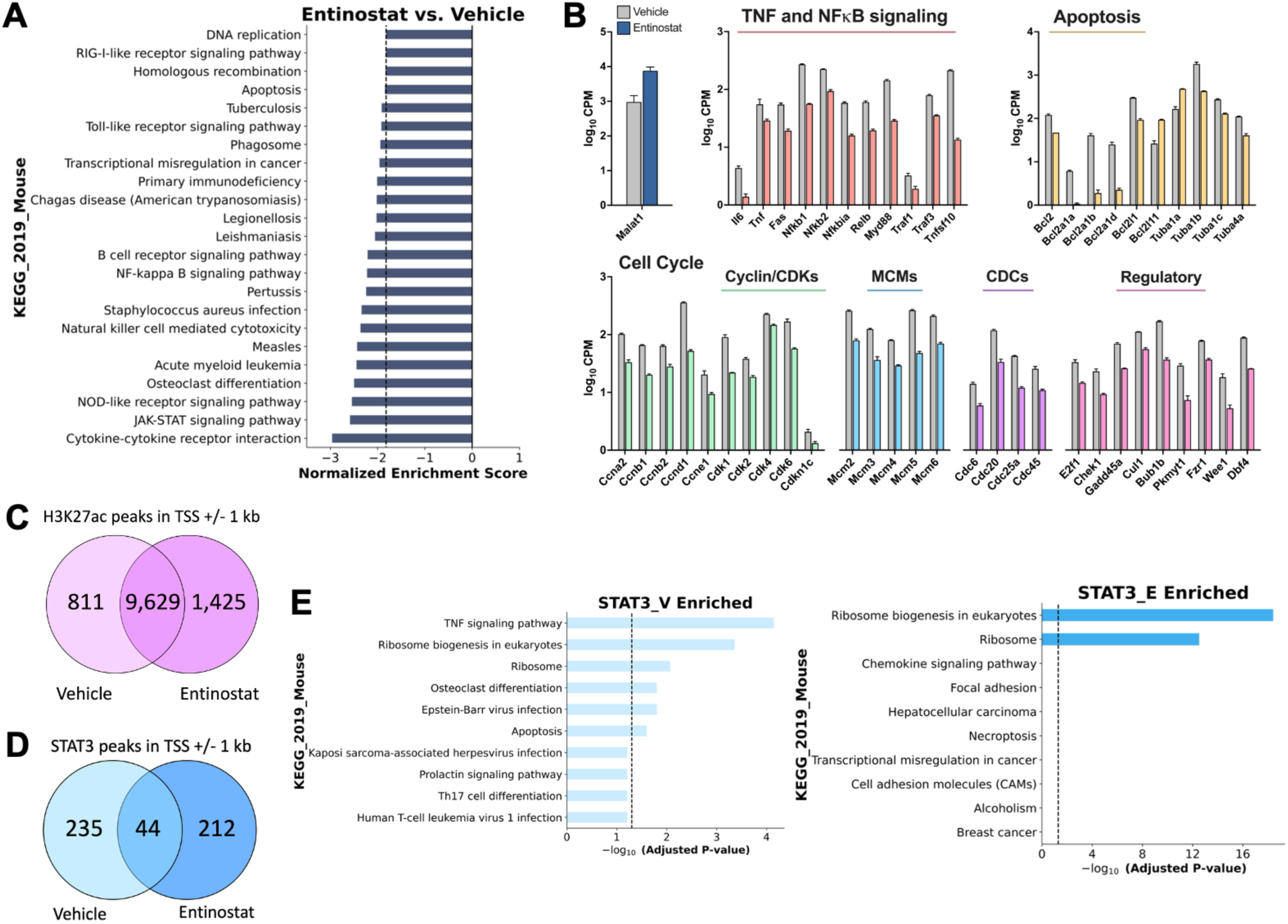
HDAC1/3 inhibition decreases apoptotic, cell cycle, and TNF signaling. **(A)** GSEA enrichment plot of significantly differentially ranked KEGG_2019_Mouse pathways from bulkRNAseq of J774M cells treated with vehicle vs. Entinostat (5 uM, 24 hours, n=3 each). **(B)** Barplots showing log10(CPM) TMM-normalized expression of *Malat1* and significantly differentially expressed genes in TNF/NF-κB signaling, apoptosis, and cell cycle pathways (KEGG_2019_Mouse) from (A) in Vehicle (gray) and Entinostat (colored) treatment conditions. Data shown are mean +/- SEM. **(C)** Venn diagram of H3K27ac enrichment peaks in promoter regions (TSS +/-1 kb) from ChIP-seq of J774M cells treated with vehicle vs. Entinostat (5 uM, 24 hours, n=3 each). TSS = transcription start site. **(D)** Venn diagram of STAT3 enrichment peaks in promoter regions (TSS +/- 1 kb) from ChIP-seq of J774M cells treated with vehicle vs. Entinostat (5 uM, 24 hours, n =3 each). TSS = transcription start site. **(E)** GSEA enrichment pathways analyses (KEGG_2019_Mouse) by enrichr method showing STAT3-vehicle (left) and –Entinostat (right) enrichment peaks in (D). Dotted line = p-adj. value cutoff of 0.05.

Using H3K27ac as a histone mark associated with actively transcribing promoter and enhancer regions, we performed chromatin immunoprecipitation sequencing (ChIP-seq) on J774M cells treated with Entinostat. Broad enrichment peak distribution analysis revealed that H3K27ac was mainly enriched around transcription start sites (TSS), followed by intronic regions and intergenic regions (**Fig. S4A**). Even though Entinostat treatment increased the total number of H3K27ac enrichment peaks (V: 55,614; E: 77,605), the percentage of H3K27ac-enriched peaks in the TSS decreased from 55.72% to 47.86%. After filtering for peaks +/- 1 kb from the TSS, we found 811 unique peaks in the vehicle condition, 1,425 unique peaks in the Entinostat condition, and 9,629 overlapping peaks (**Fig. 4C**). Pathways enrichment analysis of the condition-unique peaks by excluding the overlapping peaks did not yield many significantly enriched pathways (**Fig. S4B-G**). When comparing condition-unique peaks from differentially regulated apoptosis and cell cycling pathways identified from bulkRNA-seq, we found that Entinostat treatment decreased H3K27ac enrichment at the promoters of anti-apoptotic genes (*Bcl2a1a*, *Bcl2a1b*) and cell cycle regulatory genes (*Cdc25c, Mcm3*) (**Table S2**).

To determine which pathways may be regulated by STAT3, we performed STAT3 ChIP-seq on J774M cells treated with Entinostat. We confirmed that Entinostat treatment decreased STAT3 enrichment at its own promoter (**Fig. S5A-B**). Broad enrichment peak distribution analysis revealed that STAT3 was mainly found in intergenic regions, followed by intronic regions and transcription start sites (TSS) (**Fig. S5C**). Even though Entinostat treatment increased the total number of STAT3 enrichment peaks (V: 2,797; E: 3,913), the percentage of STAT3-enriched peaks in the TSS decreased from 14.12% to 7.16%. After filtering for peaks +/- 1 kb from the TSS, we found 235 unique peaks in the vehicle condition, 212 unique peaks in the Entinostat condition, and 44 overlapping peaks (**Fig. 4D**). Pathway enrichment analysis of the STAT3 enrichment peaks revealed STAT3 presence at the promoters of genes associated with “TNF signaling,” “Osteoclast differentiation,” “Epstein Barr virus infection,” and “Apoptosis” pathways in the vehicle condition that disappeared upon Entinostat treatment, such as *Il6, Irf1, Rela, Junb, Fos, Parp3, Gadd45b,* and *Mcl1* (**Fig. 4E, S6A-G, Table S3**). Our findings from multi-pronged sequencing suggest a role of HDAC inhibition in decreasing the expression of anti-apoptotic genes and cell cycle progression genes in MDSCs that directly affect their survival and immune-suppressive functions.

### HDAC1 inhibition increases MDSC apoptosis

HDACs have been reported to regulate processes such as cell proliferation, cell cycling, angiogenesis, and apoptosis in cancer cells (43). The STAT3 pathway has also been reported to regulate those processes in both cancer cells and MDSCs (44). Given the decreased enrichment of the apoptosis pathway from our STAT3 ChIP and the downregulation of pro-survival/anti-apoptotic Bcl-2 family genes from our bulkRNA-seq in MDSCs treated with Entinostat, we wondered if HDAC inhibition could also regulate apoptosis in MDSCs. In J774M cells, treatment with Entinostat and Stattic, a STAT3 inhibitor, for 72 hours showed visible changes in cell proliferation and cell death (**Fig. 5A**). Annexin V staining of J774M cells showed an increased percentage of apoptotic cells with Entinostat and Stattic treatment (**Fig. 5B-C**). Similarly, hMDSCs treated with Entinostat and Stattic for 48 hours showed an increased percentage of apoptotic cells (**Fig. 5D-E**). From this, we presumed that the increase in cell death would decrease cytokine production, in line with the decreased cytokine pathway enrichment from our bulkRNA-seq data. We collected supernatants from Entinostat-treated J774M cells and hMDSCs for cytokine profiling and found a decrease in the secretion of various suppression-related cytokines (**Fig. S8**), such as G-CSF, IL-6, TNF-⍺, IL-10, IL-18, PDGF-AA, APRIL, CXCL9, and CCL2, that have been reported to be involved in MDSC expansion and recruitment, regulatory T cell induction and recruitment, pro-tumoral angiogenesis, and MDSC suppression of T cells and NK cells (45–52). Our findings suggest that HDAC1 inhibition-induced apoptosis of MDSCs may be an immunomodulatory route to affect their suppressive functions.

**Figure 5:**
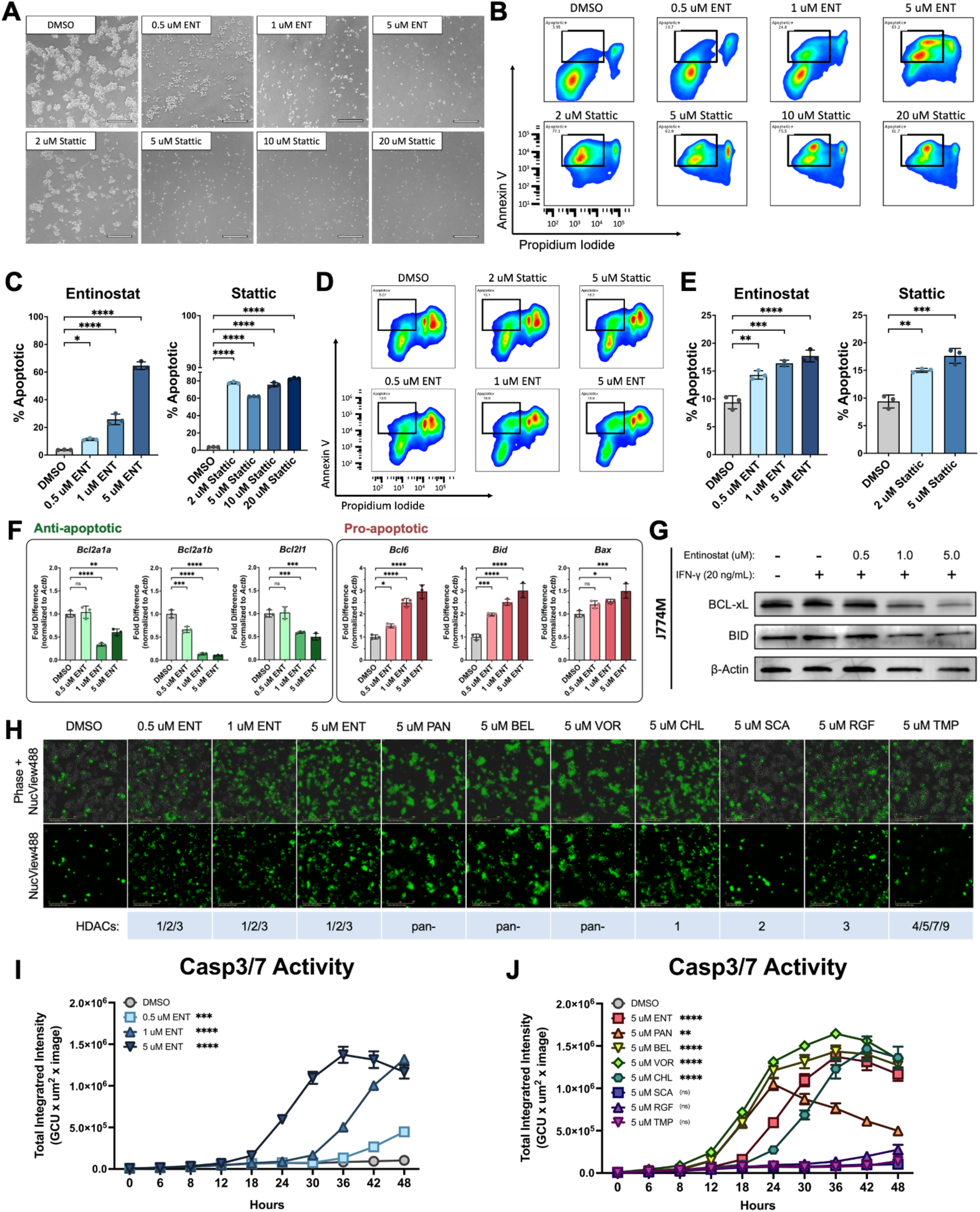
HDAC1 inhibition increases MDSC apoptosis. **(A)** Representative brightfield images of J774M cells after 72 hours of Entinostat and Stattic treatment. Scale bars = 300 um. **(B)** Representative flow cytometry plots and **(C)** percentage of apoptotic (Annexin V+, Propidium Iodide -) J774M cells after 72 hours of Entinostat and Stattic treatment (n=3). **(D)** Representative flow cytometry plots and **(E)** percentage of apoptotic hMDSCs after 48 hours of Entinostat and Stattic treatment (n=3). **(F)** Expression of anti-apoptotic genes (*Bcl2a1a, Bcl2a1b, Bcl2l1*; green) and pro-apoptotic genes (*Bcl6, Bid, Bax*; red) in J774M cells after 24 hours of Entinostat treatment (n=3). **(G)** Western blot of pro-apoptotic BCL-xL, anti-apoptotic BID, and Beta-actin in J774M cells after 24 hours of Entinostat treatment and IFN-γ stimulation (30 min, 20 ng/mL). 20 ug of total protein lysate loaded in each lane. **(H)** Representative phase + NucView488 (top) and NucView488 only (bottom) live cell images at 20x magnification of J774M cells after 48 hours of various HDAC inhibitor treatments, indicative of active caspase 3/7. Scale bars = 200 um. **(I)** Total Integrated Intensity (TII) live cell imaging timecourse quantification of NucView488 fluorescence in J774M cells after 48 hours of various concentrations of Entinostat treatment and **(J)** various HDAC inhibitor treatments, indicative of active caspase 3/7 (n=3), with TII calculated from 9 images for each replicate. Statistical significance indicated for 48-hour timepoint by One-way ANOVA. ns = not significant, * p<0.05, ** p<0.01, *** p<0.001, **** p<0.0001. ENT = Entinostat; PAN = Panobinostat; BEL = Belinostat; VOR = Vorinostat; CHL = Chlopynostat; SCA = Santacruzamate A; RGF = RGFP966; TMP = TMP269.

From multiple ChIP- and bulkRNA-seq analyses, HDAC inhibition increases MDSC apoptosis by shifting the pro-vs. anti-apoptotic signals. From this, we performed qPCR and confirmed that Entinostat treatment decreased expression of anti-apoptotic genes (*Bcl2a1a, Bcl2a1b, Bcl2l1*) and increased expression of pro-apoptotic genes (*Bcl6, Bid, Bax*) (**Fig. 5F**). At the protein level, we confirmed a decrease in expression of the anti-apoptotic protein BCL-xL encoded by *Bcl2l1*, with a less drastic change in the expression of pro-apoptotic protein BID. With this observed decrease in survival signals, we performed live cell imaging to measure the activity of executioner caspases-3 and -7. Incubation of J774M cells with NucView488 revealed that Entinostat increased the activity of caspases-3/7 in a dose- and time-dependent manner, with active caspase-3/7 detected as early as 18 hours post-treatment (**Fig. 5H-I**). Using other select HDAC inhibitors, we found that active caspase-3/7 is most robustly induced by inhibitors targeting HDAC1 (**Fig. 5H, 5J**). Our findings suggest an HDAC1-dependent shift in pro- vs. anti-apoptotic signals in MDSCs to increase caspase activity, resulting in increased apoptosis.

### HDAC1/3 inhibition increases G0/G1 cell cycle arrest in MDSCs

Cell cycling appeared as a differentially downregulated pathway from our bulkRNA-seq and the top enriched pathway from our H3K27ac ChIP-seq in MDSCs, and brightfield images of J774M cells treated with Entinostat also showed a decrease in cell proliferation. There is a long-standing link between changes in cell cycling and apoptosis (53), so we asked whether HDAC inhibition could alter cell cycling in MDSCs. We conducted propidium iodide staining to separate out cell cycle phases and found in J774M cells that treatment with Entinostat for 24 hours increased the percentage of cells in G0/G1 phase and decreased the percentage of cells in S phase (**Fig. 6A-B**). Given that the cell cycling pathway did not appear in our STAT3 ChIP, we gathered that STAT3 inhibition may not affect cell cycling in MDSCs. Indeed, Stattic treatment did not significantly decrease the percentage of J774M cells in S phase, but it did increase the percentage of cells in sub-G0/G1 phase, indicative of apoptotic cells (**Fig. 6C-D**). Using other selective HDAC inhibitors, we again found that inhibitors targeting HDAC1 significantly decreased the percentage of cells in S phase and increased those in sub-G0/G1 phase (**Fig. 6E-F**). Even though HDAC3-specific inhibition did not increase apoptosis, we did observe a similar decrease in the percentage of cells in S phase and increase in G0/G1 phase.

**Figure 6:**
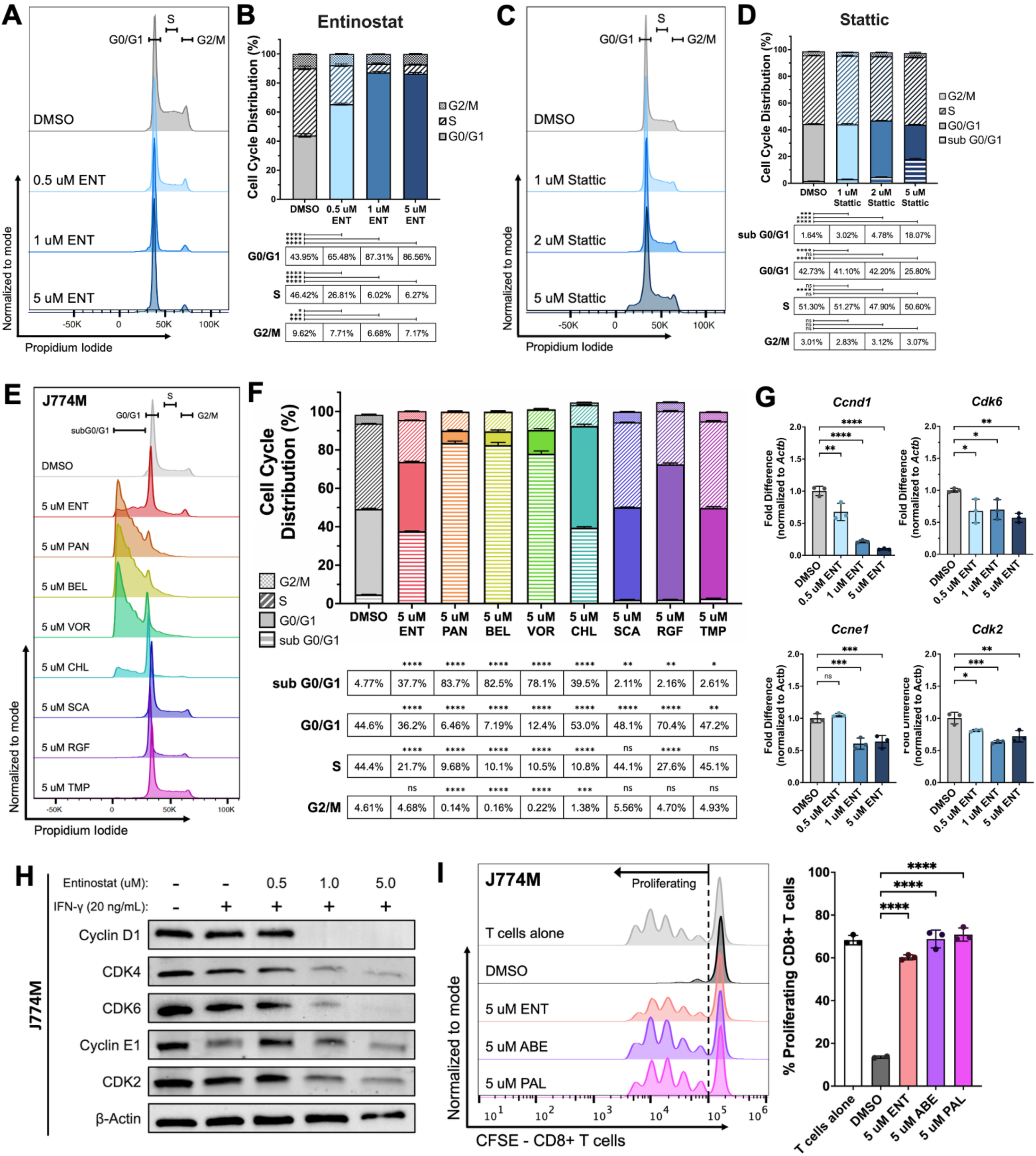
HDAC1 inhibition increases G0/G1 cell cycle arrest in MDSCs to decrease T cell suppression. **(A)** Representative flow cytometry plot and **(B)** percentage of cell cycle phase distributions in J774M cells after 24 hours of Entinostat treatment (n=3). **(C)** Representative flow cytometry plot and **(D)** percentage of cell cycle phase distributions in J774M cells after 24 hours of Stattic treatment (n=3). **(E)** Representative flow cytometry plot and **(F)** percentage of cell cycle phase distributions in J774M cells after 24 hours of various HDAC inhibitor treatments (n=3). **(G)** Expression of G1-to-S progression cell cycle genes (*Ccnd1, Cdk6, Ccne1, Cdk2*) in J774M cells after 24 hours of Entinostat treatment (n=3). **(H)** Western blot of Cyclin D1, CDK4, CDK6, Cyclin E1, CDK2, and Beta-actin in J774M cells after 24 hours of Entinostat treatment and IFN-γ stimulation (30 min, 20 ng/mL). 20 ug of total protein lysate loaded in each lane. **(I)** Representative flow cytometry plot (left) and quantification (right) of proliferating murine CD8+ T cells after 66-72 hours of co-culture with J774M cells pre-treated for 72 hours with Entinostat (ENT) and CDK4/6 inhibitors Abemaciclib (ABE) and Palbociclib (PAL) (n=3). One-way ANOVA, ns = not significant, * p<0.05, ** p<0.01, *** p<0.001, **** p<0.0001.

With our observations that HDAC inhibition increased cell cycle arrest in the G0/G1 phase and decreased progression to S phase, we wondered whether key cyclins and cyclin dependent kinase complexes that regulate G1-S transition were altered. From our bulkRNA-seq, we found that Entinostat treatment decreased expression of cyclin and cyclin dependent kinase genes *Ccnd1, Cdk6, Ccne1,* and *Cdk2* (**Table S2**) and confirmed that decrease in expression by qPCR (**Fig. 6G**). Protein analyses also showed a decrease in Cyclin D1, CDK4, CDK6, Cyclin E1, and CDK2 proteins in J774M cells with Entinostat treatment (**Fig. 6H**). From these findings, we were interested in whether FDA-approved CDK4/6 inhibitors for breast cancer could also affect the MDSC population. Pre-treatment of J774M cells with CDK4/6 inhibitors Abemaciclib (ABE) and Palbociclib (PAL) decreased their suppression of CD8+ T cells to restore proliferation (**Fig. 6I**). Our findings suggest that HDAC1/3 inhibition decreases the expression of key cyclin-CDK complexes in MDSCs that lead to G0/G1 cell cycle arrest and provide evidence supporting the use of HDAC inhibitors for specific CDK4/6 inhibition. By decreasing the expression of anti-apoptotic proteins and cyclin-CDK complex proteins, HDAC1 inhibition increases apoptosis and cell cycle arrest of MDSCs to decrease suppressive cytokine production and ultimately restore T cell proliferation (**Fig. 7**). From these findings, we propose concordant mechanisms that converge to decrease MDSC suppressive function, providing a better molecular understanding of how HDAC inhibition reprograms MDSCs.

**Figure 7:**
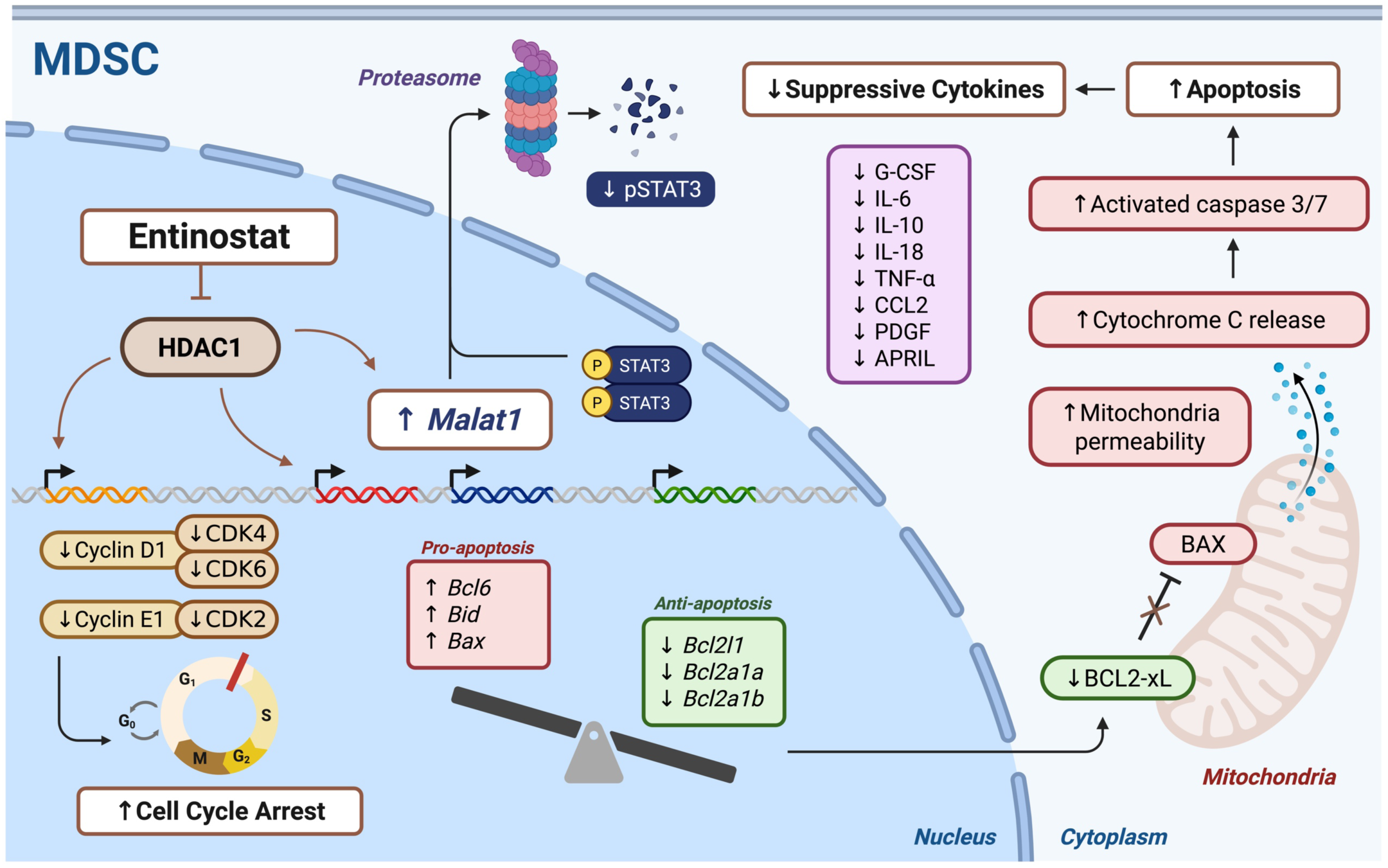
Graphic schematic of HDAC1 inhibition to decrease MDSC suppression. Entinostat’s inhibition of class I HDACs, specifically HDAC1, simultaneously influences multiple pathways of MDSC function. HDAC1 inhibition increases the expression of lncRNA *Malat1* to target phosphorylated STAT3 (pSTAT3) for ubiquitin mediated degradation by the proteasome, thereby decreasing STAT3 pathway activation. HDAC1 inhibition decreases expression of anti-apoptosis/pro-survival genes *Bcl2l1, Bcl2a1a,* and *Bcl2a1b* and increases expression of pro-apoptosis genes *Bcl6, Bid,* and *Bax*, leading to a shift in balance of pro- vs. anti-apoptosis intrinsic signaling that ultimately increases active executioner caspases 3/7 to increase apoptosis in MDSCs. HDAC1 inhibition also decreases expression of key Cyclin-CDK protein complexes to cause cell cycle arrest in the G0/G1 phase, preventing MDSC cell division and proliferation. Coupled with decreased suppressive cytokine production, HDAC1 inhibition of MDSCs to increase apoptosis and cell cycle arrest leads to a decrease in their suppression of T cells.

## DISCUSSION

Tumor-infiltrating immunosuppressive myeloid cells such as MDSCs (54) represent a significant barrier to successful immunotherapies in a wide range of cancers. Numerous studies have laid out the suppressive mechanisms by which MDSCs inhibit robust anti-tumoral T cell responses, from nutrient depletion and suppressive cytokine secretion in the TME to direct engagement of checkpoint molecules on T cells (55). However, isolated targeting of MDSCs to improve cancer response has to-date shown limited success (9). Our previous work found that ubiquitous targeting of the breast and metastatic TMEs with HDAC inhibitor Entinostat epigenetically reprograms and sensitizes the TME to ICIs, improving response by decreasing myeloid suppression (12–14,17,19). Through mathematical modeling, we have also revealed the specific immunosuppression parameters controlling response (56,57). While the use of therapeutics that simultaneously target multiple cell types and pathways more favorably produce a coordinated anti-tumoral response, the intricacies and complexities of dynamic cell-cell interactions in the TME make it challenging to tease apart mechanistic cascades. This study capitalizes on the intersectionality of statistical modeling and molecular investigations to uncover biological drivers of subtle cell states brought about by global epigenetic changes in the TME and elucidate multi-pronged mechanisms of HDAC inhibition in a key immunosuppressive cell population leading to improved response. We applied iLR to scRNA-seq data of breast-to-lung metastases-bearing mice treated with Entinostat and ICI combination therapy and identified long noncoding RNA *Malat1* as a target of HDAC inhibition in MDSCs. We confirmed in multiple MDSC models that specific HDAC1 inhibition increases *Malat1* expression to decrease STAT3 activation. Further transcriptomic and epigenomic analyses revealed changes in HDAC/STAT3-dependent apoptotic signaling and HDAC-dependent cell cycle signaling, whereby HDAC1 inhibition shifted MDSCs towards a more pro-apoptotic and cell cycle arrested state.

Our work opens various avenues for improved therapeutic combinations and molecular investigations in metastatic breast cancer. *Malat1* is a well-studied lncRNA in many cancers, with much focus on its role in cancer cell proliferation and metastasis, as well as being a potential biomarker of diagnosis and prognosis (58,59). In breast cancer, there is conflicting evidence, with some studies showing *Malat1* to be proliferation promoting and others showing *Malat1* to be metastasis suppressing (60,61). The promiscuous cellular targets of HDAC inhibitors invite future investigations into whether the mechanisms we observed in MDSCs are also observed in other immune cells and cancer cells. If *Malat1* expression in breast cancer cells, and thereby proliferation or metastatic processes, are HDAC-dependent, there may be promise in the addition of HDAC inhibitors to other therapeutics with the intent of slowing metastatic growth.

Along the same vein of therapeutic potential, one study has shown that the combination of CDK2 inhibitors with CDK4/6 inhibitors in resistant HR+/HER2- and TNBC breast cancer models provides durable tumor regression and increased survival (62). We have found in MDSCs that HDAC inhibition decreases both CDK2-Cyclin E and CDK4/6-Cyclin D complex expressions. This positions HDAC inhibition as an upstream regulator of MDSC cell cycling to be more potently immunomodulatory and cytotoxic than CDK2 or CDK4/6 inhibition alone. Additionally, our findings of HDAC inhibition causing cell cycle arrest in MDSCs corroborate clinical observations of reversible neutropenia in patients with breast cancer undergoing CDK4/6 inhibitor treatments (63). CDK4/6 inhibitors Abemaciclib, Palbociclib, and Ribociclib are FDA approved for combination with endocrine therapy in patients with HR+/HER2-breast cancer, but concurrent addition to anti-PD-1 treatment has shown to induce severe and prolonged immune-related adverse effects (irAEs) (64). Yet, the observed decrease in myeloid suppression from CDK4/6 inhibition to restore T cell proliferation in our study may prompt a revisit of this therapeutic combination to revise timing of drug administration to determine a beneficial window of maximal anti-tumor response and minimal irAEs. Collectively, our study provides insight into multi-pronged molecular mechanisms of HDAC inhibition in MDSCs and highlights the potential for our findings to be translated into clinical investigations and applications, benefiting patients with ICI-resistant metastatic breast cancer.

## MATERIALS AND METHODS

### Cell lines and culture

NT2.5-LM is a rat HER2/neu+ murine breast cancer cell line derived from NT2.5 that spontaneously metastasizes to the lungs of NeuN/FvB mice with high penetrance (20). The MDSC-like J774M cell line was kindly provided by Dr. Kebin Liu’s lab at Augusta University (65). In short, J774M cells are a CD11b+ Gr-1+ FACS sorted stable cell line from the parent J774A.1 cell line that suppress T cell proliferation. 4T1 is a triple negative breast cancer murine cell line obtained from ATCC. All cell cultures were maintained at 37°C, 5% CO_2_ in humidified incubators. NT2.5-LM and 4T1 breast cancer cells were cultured in breast cell media: RPMI-1640 (Gibco, cat. #11875-093) supplemented with 20% FBS (Gemini, cat. # 100-106), 1.5% HEPES (Gibco, cat. #15630-080), 1% L-Glutamine (Gibco, cat. #25030-081), 1% MEM non-essential amino acids (Gibco, cat. #11140-050), 1% sodium pyruvate (Sigma, cat. #S8636), 1% penicillin/streptomycin (Gibco, cat. #15140-122), and 0.2% insulin (NovoLog, cat. #NDC 0169-7501-11). J774M cells were cultured in complete media: RPMI-1640 supplemented with 10% FBS, 1.5% HEPES, 1% L-Glutamine, 1% MEM non-essential amino acids, 1% sodium pyruvate, 1% penicillin/streptomycin, 0.0004% beta-mercaptoethanol (Sigma, cat. #M3148). Cell lines were stored in liquid nitrogen and passaged at least once after thawing before experimental use, with no cell lines used after 15 passages. Only cell lines tested negative for mycoplasma were used, with testing conducted yearly.

### Animals

7-12-week-old female NeuN mice (Jackson Labs, strain #002376) were used for studies with the NT2.5-LM cell line, which were bred in-house by brother-sister mating in the animal housing facility at the University of Southern California. 7-12-week-old female BALB/c mice (Jackson Labs, strain #000651) were used for studies with the 4T1 cell line and for studies where isolated splenic T cells are co-cultured with J774M cells. All mice were housed under pathogen-free conditions and handled according to protocol approved by the USC Institutional Animal Care and Use Committee, following guidelines by the American Association of Laboratory Animal Committee policies.

### Chemicals

Chemical inhibitors used in this study: Entinostat (Syndax Pharmaceuticals), Panobinostat (MedChemExpress, cat. #HY-10224), Belinostat (MedChemExpress, cat. #HY-10225), Vorinostat (MedChemExpress, cat. #HY-10221), Trichostatin A (MedChemExpress, cat. #HY-15144), Domatinostat (MedChemExpress, cat. #HY-16012A), Chlopynostat (MedChemExpress, cat. #HY-161464), Pyroxamide (MedChemExpress, cat. #HY-13216), Santacruzamate A (MedChemExpress, cat. #HY-N0931), RGFP966 (MedChemExpress, cat. #HY-13909), TMP269 (MedChemExpress, cat. #HY-18360), SIS17 (MedChemExpress, cat. #HY-128918), Stattic (MedChemExpress, cat. #HY-13818), Abemaciclib (MedChemExpress, cat. #HY-16297A), Palbociclib (MedChemExpress, cat. #50767). Immune checkpoint inhibitors (ICIs) used in this study: anti-CTLA-4 (BioXCell cat. #BE0131) and anti-PD-1 (BioXCell cat. #BE0146).

### Human PBMC-derived MDSCs

Protocol adapted from previous studies (66,67) (**Fig. S1**). Blood from healthy donors was obtained at the University of Southern California under approved consent and IRB. Peripheral blood mononuclear cells (PBMCs) were separated by Ficoll gradient and enriched for CD33+ cells with the EasySep Human CD33+ Selection Kit II (Stem Cell, cat. #17876) per manufacturer’s instructions. These cells were plated at 5x10^5^ cells/mL in complete media for 7 days in the presence of recombinant human GM-CSF (20 ng/mL; Stem Cell, cat. #78140) and IL-6 (20 ng/mL; Stem Cell, cat. #78050.1). GM-CSF was added on days 0, 2, and 4. IL-6 was added on day 4. On day 7, the human PBMC-derived MDSCs (hMDSCs) were collected and validated for suppressive capability through MDSC-T cell co-culture proliferation assays. Following confirmation of suppressive MDSC derivation procedure, downstream experiments using inhibitors started on day 7 with re-plated hMDSCs in complete media + GM-CSF + IL-6.

### MDSC-T cell co-culture proliferation assay

*<u>Murine:</u>* CD8+ T cells were isolated from spleens of healthy 8-12-week-old female BALB/c mice with the EasySep Mouse CD8+ Isolation Kit (Stem Cell, cat. #19853) per manufacturer’s instruction. Isolated CD8+ T cells were stained with CFSE proliferation dye (ThermoFisher, cat. #C34554) at 3 uM. J774M cells were pre-treated for 72 hours with selected inhibitors and stained with Far Red proliferation dye (ThermoFisher, cat. #C34564) at 1 uM. 5x10^4^ CFSE-labeled CD8+ T cells were co-cultured with Far Red-labeled J774M cells at a 1:1-1:2 ratio (J774M:T cell) and stimulated with murine anti-CD3/CD28 dynabeads (ThermoFisher Scientific, cat. #11452D) as per manufacturer’s instructions in complete media for 66-72 hours. Afterwards, the samples were collected, stained with Live/Dead Fixable Aqua (ThermoFisher, cat. #L34965) and ran on the Attune NxT (ThermoFisher). Collected data was analyzed with FlowJo v10.10. Percent of proliferating CD8+ T cells was determined by gating on Fixable Aqua-Far Red-CFSE+ cells. For CDK4/6 inhibitor experiments, the following deviations were performed: 1x10^4^ J774M cells were plated in 96-well plates for 24 hours before replacement of media with CDK4/6 inhibitors. J774M cells were treated with CDK4/6 inhibitors for 72 hours before careful removal of supernatant containing floating dead cells, leaving live cells settled on the bottom. 5x10^4^ CFSE-labeled CD8+ T cells were then added, mixed well, and co-cultured with anti-CD3/CD28 dynabeads in complete media for 66-72 hours before collection, live/dead staining, and analysis on the Attune NxT. *<u>Human</u>*: Blood from healthy donors was obtained at the University of Southern California under approved consent and IRB. Peripheral blood mononuclear cells (PBMCs) were separated by Ficoll gradient and enriched for CD3+ cells with the EasySep Human CD3+ Selection Kit II (Stem Cell, cat. #17851) per manufacturer’s instructions. Isolated CD3+ T cells were stained with CFSE proliferation dye at 3 uM. hMDSCs derived from CD33+ enriched PBMCs as described above were pre-treated for 48 hours with selected inhibitors and stained with Far Red proliferation dye at 1 uM. 5x10^4^ CFSE-labeled CD3+ T cells were co-cultured with Far Red-labeled hMDSCs at a 1:1-1:2 ratio (hMDSC:T cell) and stimulated with human anti-CD3/CD28 dynabeads (ThermoFisher Scientific, cat. #11453D) as per manufacturer’s instructions in complete media for 66-72 hours. Afterwards, the samples were collected, stained with Live/Dead Fixable Aqua (ThermoFisher, cat. #11131D) and ran on the Attune NxT (ThermoFisher). Collected data was analyzed with FlowJo v10.10. Percent of proliferating CD3+ T cells was determined by gating on Fixable Aqua-Far Red-CFSE+ cells.

### Murine studies

For single cell RNA sequencing (scRNA-seq) of NT2.5-LM metastatic lung tumors and isolation of NT2.5-LM G-MDSCs, detailed procedures as previously described (17,19,20). For scRNA-seq, 1x10^5^ NT2.5-LM cells were injected in the mammary fat pad of 7-12-week-old female NeuN mice, and after surgical resection of the breast tumor on day 12 post-injection, mice began treatments on day 17 post-injection: 1) Vehicle: 0.5% methylcellulose +/- isotype antibodies in PBS; 2) Entinostat: 5 mg/kg, 5x/week, oral gavage; 3) ICIs: anti-CTLA-4 + anti-PD-1, 100 ug/dose, 2x/week, intraperitoneal (i.p.) injection; 4) Entinostat + ICIs: Entinostat treatment started one day prior to ICIs, same dosage and frequency. For isolation of MDSCs, the same conditions as described above, with only vehicle and Entinostat treatment groups. Treatments lasted for 3 weeks before collection of lungs for dissociation on the gentleMACS Octo Dissociator (Miltenyi) with the mouse tumor dissociation kit (Miltenyi, cat. #130-096-730), following manufacturer’s instruction. Dissociated tumors were passed through a 70 um strainer into single-cell suspensions. Red blood cells were lysed with ACK lysis buffer (Gibco, cat. # A1049201) for 5 minutes at RT. For PrimeFlow analyses in 4T1 metastatic lung tumors, 1x10^4^ 4T1 cells were injected intravenously in the lateral tail vein of 7-12-week-old female Balb/c mice, with treatments beginning on day 4 post-injection: 1) Vehicle: 0.5% methylcellulose +/- isotype antibodies in PBS; 2) Entinostat: 5 mg/kg, 5x/week, oral gavage; 3) Entinostat + ICIs: Entinostat treatment started one day prior to ICIs, anti-CTLA-4 + anti-PD-1, 100 ug/dose, 2x/week, intraperitoneal (i.p.) injection. Treatments lasted for 3 weeks, with tissue collection and processing procedures the same as described above. PrimeFlow procedures described below in corresponding section.

### Single cell RNA sequencing (scRNA-seq) and iterative logistic regression (iLR)

scRNA-seq was previously performed on lung metastases collected from the NT2.5-LM model of breast cancer, in which mice were treated with vehicle (V) or Entinostat + anti-PD-1 + anti-CTLA-4 (EPC) (19). Cells were clustered together using Louvain clustering, and 8 main clusters were identified with resolution parameter 0.1: cancer cells, mature myeloid, MDSCs, T cells, B cells, NK cells, lung endothelial cells, and lipofibroblasts. Within the MDSC cluster, subclusters of G-MDSCs and M-MDSCs were identified (19). Iterative logistic regression (iLR) was applied to the main MDSC cluster to compare marker genes between V and EPC treatment groups (21). 10 rounds of iLR with genes appearing more than 5 times resulted in the final iLR-identified gene set, with ribosomal genes and -Rik genes removed.

### Murine ex vivo MDSCs

G-MDSCs were isolated from single-cell suspensions of NT2.5-LM lung metastases after 3 weeks of vehicle vs. Entinostat treatment (as described above) using a Ly-6G+ magnetic bead pulldown from the MDSC isolation kit (Miltenyi, cat. #130-094-538), following manufacturer’s instruction. M-MDSCs were isolated from flow-throughs using a Gr-1+ magnetic bead pulldown from the same kit, following manufacturer’s instruction.

### RNA isolation and qPCR analysis

J774M cells and hMDSCs were treated with inhibitors for 24 hours and RNA was extracted using the Quick-RNA Miniprep kit (Zymo Research, cat. #R1055) following manufacturer’s instruction. cDNA synthesis was conducted on 500-1000 ng of RNA with Maxima First Strand cDNA Synthesis Kit for RT-qPCR (ThermoFisher, cat. #K1642). 5 ng of cDNA per triplicate per sample were used for qPCR using PowerUp SYBR Green Master Mix for qPCR (ThermoFisher, cat. #A25778). Primers were designed and ordered from IDT (**Table S5**).

### PrimeFlow

Lungs with metastases from 4T1 mice were collected and dissociated as described above. 1x10^6^ cells per sample were plated in a 96-well plate and stained following manufacturer’s protocol (ThermoFisher, cat. #88-18005-210) with the following alterations: viability staining was performed with Live Dead Blue at a 1:1000 dilution for 30 min at 4°C, followed by incubation with CD16/CD32 Fc receptor block (Biolegend, cat. #101302) at a 1:50 dilution for 15 minutes at room temperature (RT) before surface antibody staining (**Table S4**) for 1 hour at 4°C. Cells were fixed and permeabilized with PrimeFlow kit reagents, followed by another by incubation with CD16/CD32 Fc receptor block (Biolegend, cat. #101302) at a 1:50 dilution for 15 minutes at room temperature (RT) before intracellular antibody staining (**Table S4**) for 1 hour at 4°C. Cells were then fixed for RNA and incubated with anti-mouse *Malat1*-AlexaFluor488 probes (ThermoFisher, Type 4 RNA Probe set) following manufacturer’s protocol. Stained samples were run on the Cytek Aurora. Collected data was analyzed with SpectroFlo and FlowJo v10.10.

### Western blots

G-MDSCs isolated from NT2.5-LM lung metastasis-bearing NeuN mice (as described above) were plated at a maximum of 1x10^6^ cells/mL in complete media and stimulated with LPS (1 ug/mL, Sigma, cat. #L2630-10MG) for 2 hours at 37°C. J774M cells were plated in complete media and treated with inhibitors for 24 hours and stimulated with IFN-γ (20 ng/mL, Stem Cell, cat. #78021) for 30 minutes at 37°C. hMDSCs were plated in complete media supplemented with GM-CSF and IL-6 (as described above) and treated with inhibitors for 24 hours at 37°C. All cells were lysed with RIPA buffer (ThermoFisher, cat. #89900) supplemented with phosphatase/protease inhibitor (Cell Signaling, cat. #5872S). Whole cell lysate protein concentration was determined using BCA Protein Assay Reagent (Pierce, cat. #23225). 20-30 ug of J774M protein lysate and 50 ug of NT2.5-LM G-MDSC and hMDSC protein lysate were loaded on a 4-15% Mini-PROTEAN TGX gel (Biorad, cat. #4561083) and transferred to PVDF membranes (Millipore, cat. #IEVH00005). Membranes were blocked with 5% non-fat dry milk in TBS plus 0.1% Tween 20 (TBST) for at least 1 hour at room temperature (RT) with gentle shaking. Membranes were then incubated with primary antibodies in 5% non-fat dry milk in TBST at specified concentrations (**Table S4**) overnight at 4°C with gentle shaking. The next day, membranes were washed with TBST and incubated with secondary antibodies in 5% non-fat dry milk in TBST at specified concentrations (**Table S4**) for at least 1 hour at RT with gentle shaking. Membranes were washed with TBST, and proteins were visualized with SuperSignal West Pico PLUS Chemiluminescent Substrate (ThermoFisher, cat. #34580) for HRP-conjugated secondary antibodies. For DyLight-800 conjugated secondary antibodies, proteins were directly visualized on the LICOR Odyssey CLx Imager. Membranes were stripped up to three times using Restore PLUS Western Blot Stripping Buffer (ThermoFisher, cat. #46430) and re-probed for other protein targets. Quantification of pSTAT3, STAT3, and β-actin of ex vivo NT2.5-LM G-MDSCs conducted with ImageJ, with adjusted relative densities normalized to β-actin.

### J774M STAT3-responsive GFP-reporter generation

J774M STAT3-responsive GFP-reporter (STAT3r-GFP) was generated by lentivirus transduction with Lenti-X 293T cells (Takara Bio, cat. #632180) using the “STAT3-GFP” plasmid, a gift from Michael Lewis (Addgene, plasmid #110495) (68). In brief, Lenti-X cells were seeded at 5x10^5^ cells/well in a 6-well plate one day prior to transfection. Transfection of psPAX2 (Addgene, plasmid #12260), pMD2.G (Addgene, plasmid #12259), and STAT3-GFP plasmids were carried out with X-tremeGENE HP DNA transfection reagent (Roche, cat. #06-366-244-001) following manufacturers protocol, using Opti-MEM (ThermoFisher, cat. #31985070) for the transfection mix. Virus was collected 72 hours post-transfection and transduced into J774M cells by spinfection at 600 xg for 2 hours at 37°C in a 6-well plate with 10 ug/mL of polybrene (Sigma, cat. #TR-1003-G). After 3 days of rest, STAT3-GFP construct-expressing J744M cells were selected with 8 ug/mL of puromycin (Sigma, cat. #P8833-10MG) for 3 days. After 3 days of rest, J774M STAT3r-GFP cells were stimulated for 5 hours with IFN-γ (20 ng/mL, Stem Cell, cat. #78021) before FACS sorting for GFP+ cells. GFP+ cells were returned to culture and passaged for 3 passages before confirmation of GFP signal returning to baseline by comparing with un-stimulated/non-FACS sorted cells.

### Bulk RNA sequencing (bulkRNA-seq)

J774M cells were treated for 24 hours with vehicle or Entinostat (5 uM) and stimulated with IFN-γ (20 ng/mL) in triplicates. After washing twice in PBS, RNA was collected (Zymo Research, cat. #R1055), following manufacturer’s instruction. mRNA library with poly A enrichment was prepared and sequenced by Novogene using NovaSeq X Plus to generate approximately 25-35 million reads per sample, with average Phred quality score > 39.3. Paired-end reads were processed on PartekFlow v12.3.0 by trimming adapters and trimming reads for Phred quality score > 35.0 before alignment to the mm39 transcriptome using STAR v2.7.8a with default settings. Genes with counts <= 10.0 were removed. Counts were normalized using Median Ratio for DESeq2 analysis before differential gene analysis was conducted using DESeq2. Resulting differentially expressed gene set was further filtered and sub-setted using Python v3 for p-adj. < 0.05 and log2FoldChange >1 and <-1. Gene set enrichment analyses were performed with GSEA and Enrichr over-representation analysis methods using GSEApy v1.1

### Chromatin immunoprecipitation qPCR (ChIP-qPCR) and sequencing (ChIP-seq)

J774M cells were treated for 24 hours with vehicle or Entinostat (5 uM) and stimulated with IFN-γ (20 ng/mL). After washing twice in PBS, 2x10^7^ cells were fixed with 1% formaldehyde for 8 minutes at RT with rotation. Fixation was stopped by adding glycine to a final concentration of 0.125 M. Cell pellets were flash frozen in liquid nitrogen before proceeding to chromatin sonication. Samples were resuspended in 5 mL of freshly prepared, ice-cold Cell Lysis Buffer (5 mM PIPES [pH 8.0], 85 mM KCl, 1% NP-40, 1x protease inhibitors) for 15 minutes on ice to allow for cell lysis. Nuclear pellets were resuspended in freshly prepared, ice-cold Nuclei Lysis Buffer (50 mM Tris-HCl [pH 8.1], 10 mM EDTA, 1% SDS, 1x protease inhibitors) at a calculated concentration of 1.5x10^7^ original cells/mL for 30 minutes on ice. Samples were additionally flash frozen in liquid nitrogen to lyse nuclei, thawed on ice, and sonicated on the Diagenode Bioruptor Pico sonicator until optimal nucleosome range of 200-500 bp was achieved, as confirmed on a 1.5% agarose gel. 50 ug of chromatin in IP Dilution Buffer (50 mM Tris [pH 7.4], 150 mM NaCl, 1% NP-40, 0.25% Deoxycholic acid, 1mM EDTA, 1x protease inhibitors) was incubated with 10 uL of STAT3 (Cell Signaling, cat. #12640) and H3K27ac (Cell Signaling, cat. #8173) antibodies overnight at 4°C with rotation, in triplicates. Immunoprecipitation was carried out the following day with pre-cleared Protein A/G magnetic beads (Invitrogen, cat. #10002D and #10004D) for 2 hours at 4°C with rotation. Samples were magnetically separated, washed twice with IP Wash Buffer 1 (50 mM Tris [pH 7.4], 150 mM NaCl, 1% NP-40, 0.25% Deoxycholic acid, 1 mM EDTA) and twice with IP Wash Buffer 2 (100 mM Tris-HCl [pH 9.0], 500 mM LiCl, 1% NP-40, 1% Deoxycholic acid). Antibody-chromatin complexes were eluted by vortexing for 30 min at 37°C in IP Elution Buffer (50 mM NaHCO_3_, 1% SDS), separating magnetic beads, and collecting supernatant. Along with 1% input, samples were incubated with 0.6 M NaCl overnight at 67°C to reverse formaldehyde cross-links. The following day, DNA was purified and eluted for ChIP-qPCR and ChIP-seq. For STAT3-ChIP-qPCR, primers were designed in the promoter regions of *Stat3* (reported to be bound by STAT3) and *Gapdh* (reported to not be bound by STAT3) (**Table S5**). ChIP-seq was performed by Novogene using NovaSeq PE150 for pair-end sequencing to generate 30+ million reads per sample, with 90% of reads for each sample having a Phred quality score of > 30.0). Paired-end reads were processed on PartekFlow v12.3.0 by trimming adapters and trimming reads for Phred quality score > 20 from the 3’ end, resulting in an average Phred quality score > 35.21. Reads were aligned to the mm10 genome using Bowtie 2.2.5 with default settings. Peaks were called using MACS 2.1.1 and evaluated for distribution across genomic features. Peaks were filtered to be +/- 1kb from the transcription start site (TSS) for pathway enrichment analyses.

### Annexin V apoptosis assay

J774M cells treated with inhibitors for 72 hours and hMDSCs treated with inhibitors for 48 hours were collected, washed twice with PBS, and stained with Annexin V-FITC and Propidium Iodide (PI) using the Dead Cell Apoptosis Kits with Annexin V (ThermoFisher, cat. #V13242) following manufacturer’s instruction. Samples were kept at 4°C until they were run on the Attune NxT (ThermoFisher). Collected data was analyzed with FlowJo v10.10. Apoptotic cells were gated on Annexin V-FITC+ PI-cells.

### Live Cell Imaging

J774M cells were plated at 5x10^4^ cells/well in 96-well black imaging microplates and allowed to settle before addition of NucView488 (Biotium, cat. #30029) at a 1:400 volumetric ratio and various HDAC inhibitors, in triplicates. Live cell imaging was conducted using the Incucyte SX5 Live-Cell analysis system (Sartorius) at normal cell culture conditions, with timecourse green fluorescence and phase images taken at 20x objective spanning 9 regions in each well over 48 hours. Total integrated intensity (TII) of green fluorescence was calculated for each well: (GCU: green fluorescence intensity) x um^2^ x image.

### Multi-plex ELISAs

*<u>Murine:</u>* Supernatants were collected from J774M cells treated with inhibitors and stimulated with IFN-γ (20 ng/mL, Stem Cell, cat. #78021) for 72 hours. Multi-plexing analysis with the Mouse Cytokine/Chemokine 32-Plex Discovery Assay Array (MD32) was performed using the Luminex™ 200 system (Luminex, Austin, TX, USA) at Eve Technologies (Calgary, Alberta). The 32-plex consisted of Eotaxin-1, G-CSF, GM-CSF, IFN-γ, IL-1α, IL-1β, IL-2, IL-3, IL-4, IL-5, IL-6, IL-7, IL-9, IL-10, IL-12p40, IL-12p70, IL-13, IL-15, IL-17A, IP-10/CXCL10, KC/GRO/CINC-1/CXCL1, LIF, LIX/CXCL5, MCP-1/CCL2, M-CSF, MIG/CXCL9, MIP-1α/CCL3, MIP-1β/CCL4, MIP-2/CXCL2, RANTES/CCL5, TNFα, VEGF-A. Assay sensitivities of these markers range from 0.3-30.6 pg/mL. A cubic spline and 5-paramter logistic regression were used when optimizing the standard. Regression analysis was performed utilizing the Bio-Plex Manager software. *<u>Human</u>*: Supernatants were collected from hMDSCs treated with inhibitors for 72 hours. Multi-plexing analysis was performed by Eve Technologies Corporation (Calgary, Alberta, Canada) using the Luminex® 200™ system (Luminex Corporation/DiaSorin, Saluggia, Italy) with Bio-Plex Manager™ software (Bio-Rad Laboratories Inc., Hercules, California, United USA). The Luminex® xMAP® technology was used to quantitatively and simultaneously detect ninety-six human cytokines, chemokines and growth factors simultaneously. Ninety-six markers were measured in the samples using two separate Eve Technologies’ panels, as per the manufacturer’s instructions for use: 1) Human Cytokine/Chemokine Panel A 48-Plex Discovery Assay® Array (HD48A) (MILLIPLEX® Human Cytokine/Chemokine/Growth Factor Panel A Magnetic Bead Panel Cat. # HCYTA-60K, MilliporeSigma, Burlington, Massachusetts, USA); 2) Human Cytokine/Chemokine Panel B 48-Plex Discovery Assay® Array (HD48B) (MILLIPLEX® Human Cytokine/Chemokine/Growth Factor Panel B Magnetic Bead Panel Cat. # HCYTB-60K, MilliporeSigma, Burlington, Massachusetts, USA). The 96-plex consisted of sCD40L, sCD137, 6Ckine/CCL21, APRIL, BAFF/BLyS, BCA-1/BLC/CXCL13, CCL28, CTACK/CCL27, CXCL16, EGF, ENA-78/CXCL5, Eotaxin/CCL11, Eotaxin-2/CCL24, Eotaxin-3/CCL26, FGF-2, FLT-3 Ligand, Fractalkine/CXCL1, GCP-2/CXCL6, G-CSF/CSF-3, GM-CSF, Granzyme A, Granzyme B, GROα/CXCL1/KC/CINC-1, HMGB1/HMG1, I-309/CCL1, IFN-α2, IFN-γ, IFN-β, IFN-ω, IL-1α, IL-1β, IL-1RA, IL-2, IL-3, IL-4, IL-5, IL-6, IL-7, IL-8/CXCL8, IL-9, IL-10, IL-11, IL-12(p40), IL-12(p70), IL-13, IL-15, IL-16, IL-17A/CTLA-8, IL-17E/IL-25, IL-17F, IL-18, IL-20, IL-21, IL-22, IL-23, IL-24, IL-27, IL-28A/IFN-λ2, IL-29/IFN-λ1, IL-31, IL-33, IL-34, IL-35, IP-10/CXCL10, I-TAC/CXCL11, LIF, Lymphotactin/XCL1, MCP-1/CCL2, MCP-2/CCL8, MCP-3/CCL7, MCP-4/CCL13, M-CSF, MDC/CCL22, MIG/CXCL9, MIP-1α/CCL3, MIP-1β/CCL4, MIP-1Ᵹ/CCL15, MIP-3α/CCL20, MIP-3β/CCL19, MPIF-1/CCL23, PDGF-AA, PDGF-AB/BB, Perforin, RANTES/CCL5, SCF, SDF-1α+β/CXCL12, sFAS, sFASL, TARC/CCL17, TGF-α, Thrombopoietin (TPO), TNF-α, TNF-β, TRAIL, TSLP and VEGF-A. Assay sensitivities of these markers range from 0.05 – 100 pg/mL. Individual analyte sensitivity values are available in the MilliporeSigma MILLIPLEX® protocols.

### Propidium iodide cell cycle distribution assay

J774M cells treated with inhibitors for 24 hours were collected, washed twice with ice-cold PBS, fixed with ice-cold 70% EtOH (added dropwise with vortexing) for 30 minutes at 4°C, washed twice with PBS, and resuspended in PI staining solution (50 ug/mL, Sigma, cat. #P4170-25MG) with RNAse A (0.5 ug/mL, Cell Signaling, cat. #7013S) for at least 1 hour at 4°C. Samples were kept at 4°C until they were run on the Attune NxT (ThermoFisher). Collected data was analyzed with FlowJo v10.10 using the Cell Cycle platform and the Watson subtractive model of cell cycle distribution, with G1 and G2 histogram widths constrained to be the same.

### Statistical Analyses

Statistical analyses for sequencing data mentioned in the corresponding sections. Statistical analyses for western blots, qPCRs, flow cytometry analyses, and ELISAs were performed using GraphPad Prism 8 software. To assess statistical significance, One-way ANOVAs and Two-way ANOVAs were used when appropriate, with post hoc analyses conducted using Tukey’s multiple comparison tests. For murine flow cytometry studies, Kruskal-Wallis tests with Dunn’s correction for post hoc analyses were used. Histograms show individual data points with mean +/- SD (Standard Deviation), with the threshold of significance set at p < 0.05.

## Supplementary Materials

Fig. S1: Human PBMC-MDSC and T cell co-culture derivation schematic

Fig. S2: J774M STAT3r-GFP reporter

Fig. S3: GSEA pathways enrichment from J774M bulkRNA-seq

Fig. S4: H3K27ac ChIP-seq peak distributions and GSEA pathways enrichment of treatment specific peaks

Fig. S5: STAT3 ChIP-qPCR and ChIP-seq peak distributions

Fig. S6: STAT3 ChIP-seq GSEA pathways enrichment of treatment specific overall peaks

Fig. S7: Class I HDAC inhibition decreases suppressive cytokine secretion

Table S1: J774M BulkRNA-seq GSEA and DEGs

Table S2: J774M H3K27ac ChIP-seq GSEA

Table S3: J774M STAT3 ChIP-seq GSEA

Table S4: Western blot and flow cytometry antibodies

Table S5: qPCR primer sequences

## Acknowledgements

We would like to acknowledge the core facilities at USC Norris Comprehensive Cancer Center, including the Molecular Genomics Core, Flow Cytometry and Immune Monitoring Core, and Stem Cell Flow Cytometry Core. We would also like to thank Min Yu and Des Mecenas for their advice throughout the course of the project, Jasmine Martinez for her help with molecular techniques, and Bernadette Masinsin and Junji Watanabe for their help with core services. We would like to thank Syndax for providing Entinostat.

## Author contributions

Conceptualization: AGB, ERT

Methodology: AGB, YL, EG, BA, MI, AHL, NGK, JK, JN, ALM, ERT

Investigation: AGB, YL, EG, BA, MI, AHL, NGK, MBJ, CP, JK, LK, SKZ, KA, ALM, ERT

Visualization: AGB, YL, ALM, ERT

Funding acquisition: ALM, ERT

Project administration: ERT

Supervision: ERT

Writing – original draft: AGB, ERT

Writing – review & editing: AGB, YL, EG, BA, MI, AHL, NGK, MBJ, CP, JK, LK, SKZ, KA, SJP, JN, ALM, ERT

## Data and materials availability

Bulk RNA-sequencing and ChIP-sequencing data of Entinostat-treated J774M cell lines will be deposited in the Gene Expression Omnibus (GEO) at accession numbers GSE325737 and GSE325738, respectively. scRNA-sequencing of NT2.5-LM lung metastases are accessible at GEO GSE303155.

## Ethics approval and consent to participate

All animal studies were approved by the Institutional Review Board of USC.

