## Supplemental Figures for "A class act: HDAC1-*Malat1* regulates MDSC apoptosis and cell cycling to decrease suppression of T cells"

**Supplemental Figure 1**

**A**

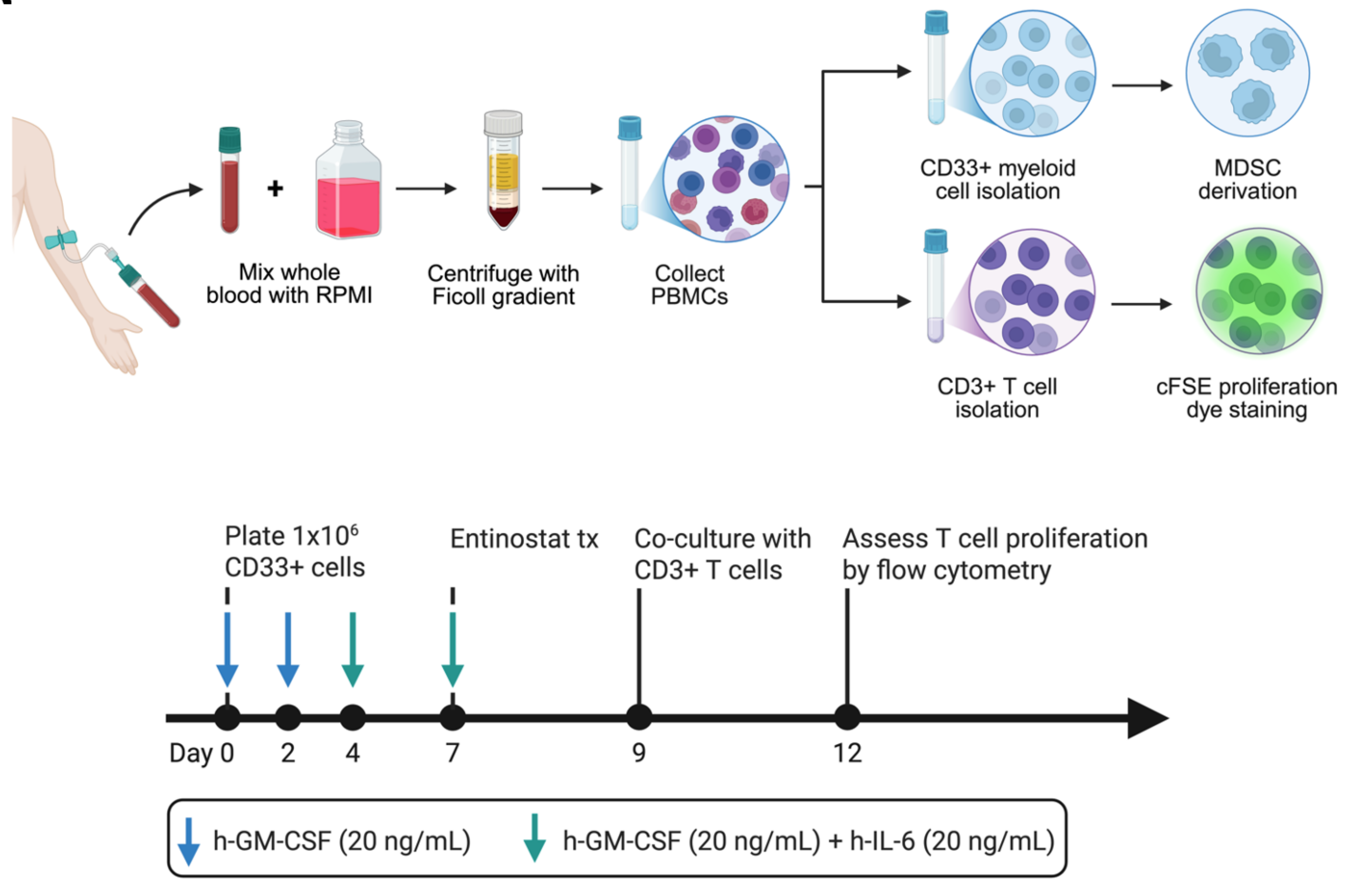


**Figure S1: Human PBMC-MDSC and T cell co-culture derivation schematic**. **(A)** Schematic of human PBMC-MDSC (hMDSC) derivation. PBMCs were isolated by Ficoll gradient centrifugation of whole blood collected from healthy donors. PBMCs were enriched for CD33+ myeloid cells before plating at $5x10^{5}$ cells/mL in 2 mL of MDSC-derivation media. MDSCs were derived after 7 days with the addition of human recombinant GM-CSF (20 ng/mL; Days 0, 2, 4), and human recombinant IL-6 (20 ng/mL; Day 4) to CD33+ myeloid cells. For T cell proliferation co-culture assays, hMDSCs were pre-treated with Entinostat on day 7 for 48 hours before collection and co-culture with CD3+ T cells. After 66-72 hours of co-culture, proliferating CD3+ T cells were assessed by flow cytometry.

**Supplemental Figure 2**

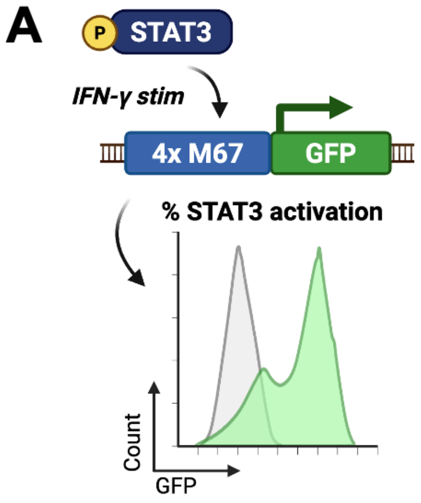


**Figure S2: J774M STAT3-responsive GFP reporter**. **(A)** Schematic of STAT3-responsive GFP reporter construct in J774M cells. Plasmid with 4 repeats of the M67 STAT3-specific DNA binding sequence is inserted in front of the GFP coding sequence. Upon immune stimulation (e.g. with IFN-gamma), STAT3 is phosphorylated and activated, inducing expression of GFP.

**Supplemental Figure 3**

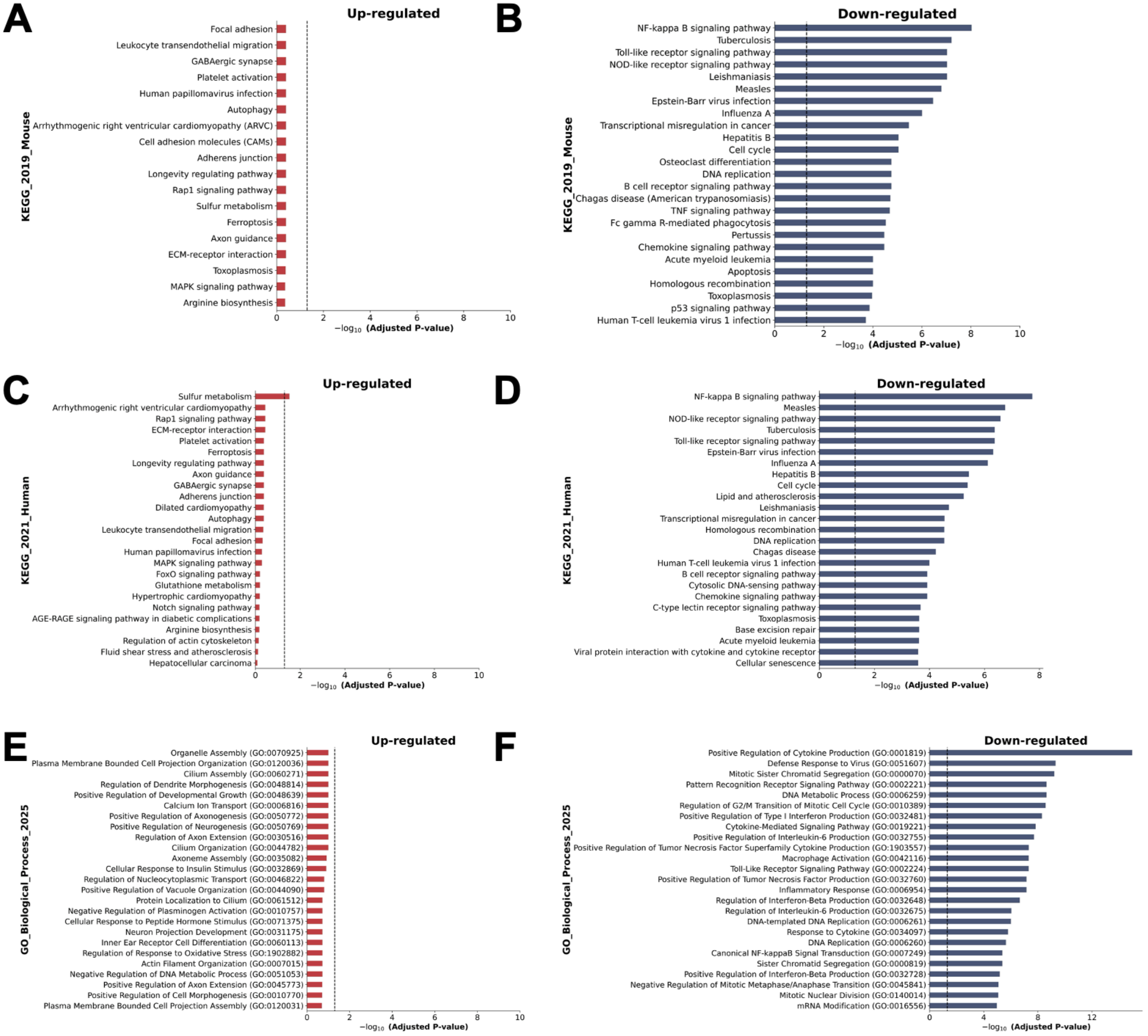


**Figure S3: GSEA pathways enrichment from J774M bulkRNA-seq**. **(A)** KEGG_2019_Mouse GSEA enrichment barplot by enrichr method showing up-regulated and **(B)** down-regulated pathways from bulkRNAseq of J774M cells treated with Entinostat. **(C)** KEGG_2021_Human GSEA enrichment barplot by enrichr method showing up-regulated and **(D)** down-regulated pathways from bulkRNAseq of J774M cells treated with Entinostat. **(E)** GO_Biological_Process_2025 GSEA enrichment barplot by enrichr method showing up-regulated and **(F)** down-regulated pathways from bulkRNAseq of J774M cells treated with Entinostat. Dotted line = p-adj. value cutoff of 0.05.

**Supplemental Figure 4**

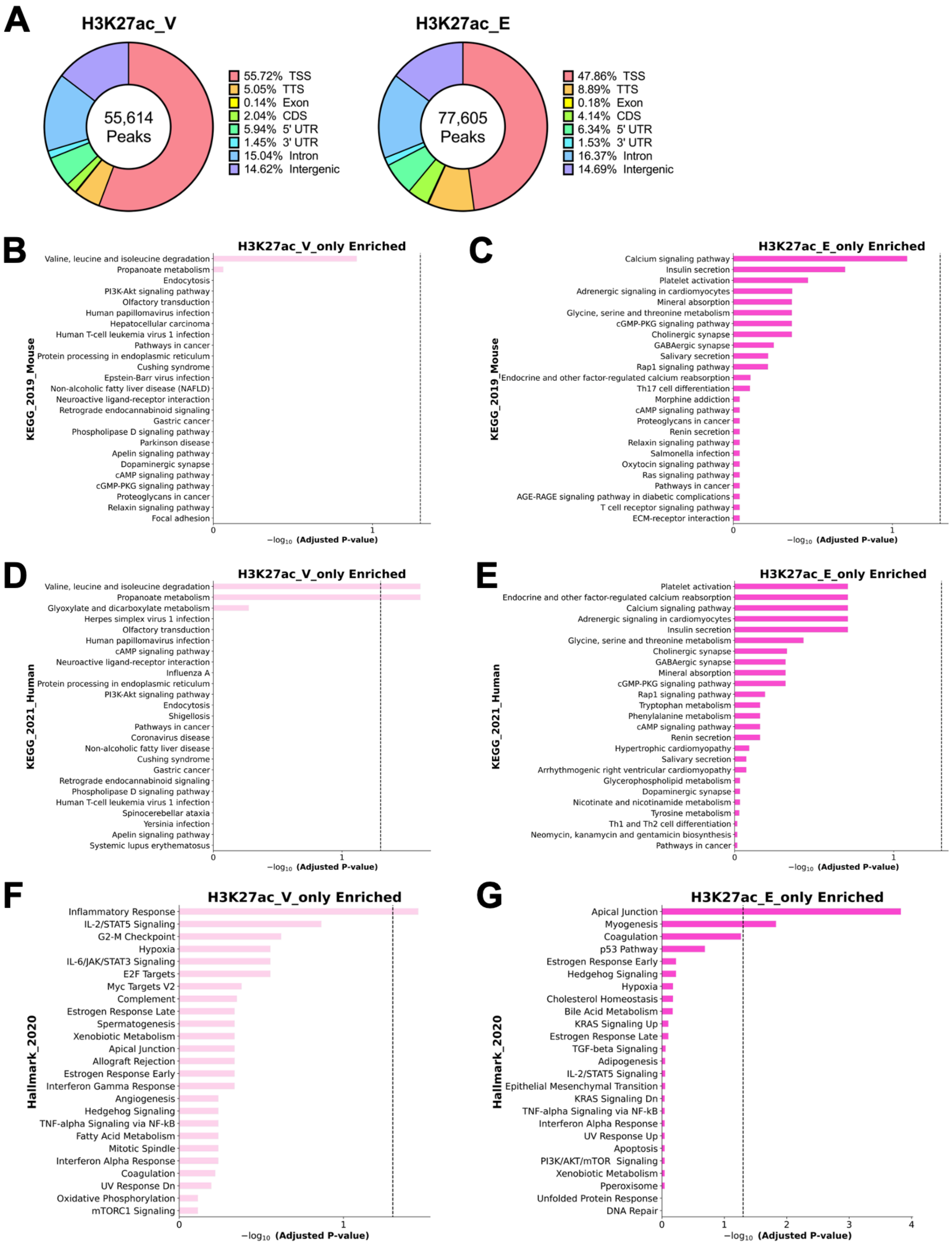

**Figure S4: H3K27ac ChIP-seq peak distributions and GSEA pathways enrichment of treatment specific peaks**. **(A)** H3K27ac-IP ChIP-seq peak distribution in J774M cells between vehicle and Entinostat treatment groups. TSS = transcription start site, TTS = transcription termination site, CDS = coding DNA sequence, 5' UTR = 5 prime untranslated region, 3' UTR = 3 prime untranslated region. **(B)** KEGG_2019_Mouse GSEA enrichment barplot by enrichr method showing H3K27ac-vehicle unique-peak and **(C)** H3K27ac-Entinostat unique-peak enriched pathways from H3K27ac ChIP-seq of J774M cells. **(D)** KEGG_2021_Human GSEA enrichment barplot by enrichr method showing H3K27ac-vehicle unique-peak and **(E)** H3K27ac-Entinostat unique-peak enriched pathways from H3K27ac ChIP-seq of J774M cells. **(F)** Hallmark_2020 GSEA enrichment barplot by enrichr method showing H3K27ac-vehicle unique-peak and **(G)** H3K27ac-Entinostat unique-peak enriched pathways from H3K27ac ChIP-seq of J774M cells. Dotted line = p-adj. value cutoff of 0.05.

**Supplemental Figure 5**

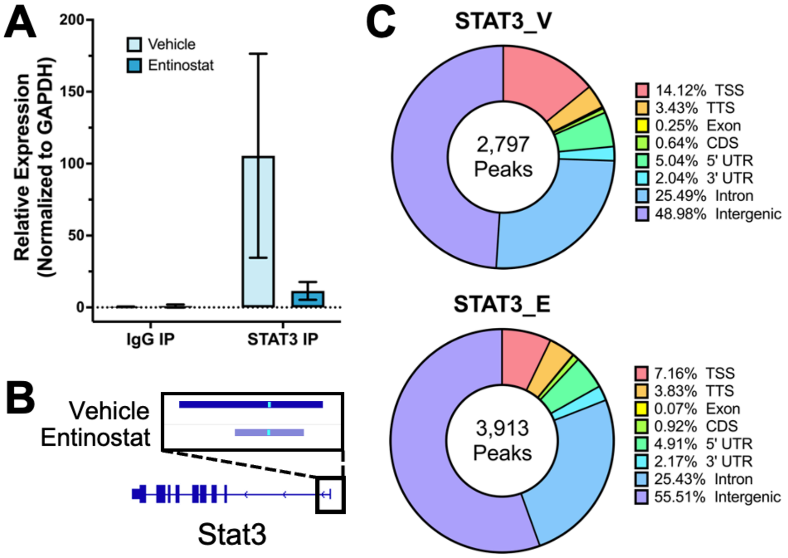


**Figure S5: STAT3 ChIP-qPCR and ChIP-seq peak distributions**. **(A)** ChIP-qPCR of IgG- vs. STAT3-immunoprecipitated genomic DNA at *Stat3* promoter, normalized to *Gapdh* promoter, in J774M cells after 24 hours of Entinostat treatment and IFN-γ stimulation (20 ng/mL). **(B)** STAT3-IP ChIP-seq NarrowPeak view of *Stat3* promoter in J774M cells between vehicle and Entinostat treatment groups, normalized across triplicates. **(C)** STAT3-IP ChIP-seq peak distribution in J774M cells between vehicle and Entinostat treatment groups. TSS = transcription start site, TTS = transcription termination site, CDS = coding DNA sequence, 5' UTR = 5 prime untranslated region, 3' UTR = 3 prime untranslated region.

### Supplemental Figure 6

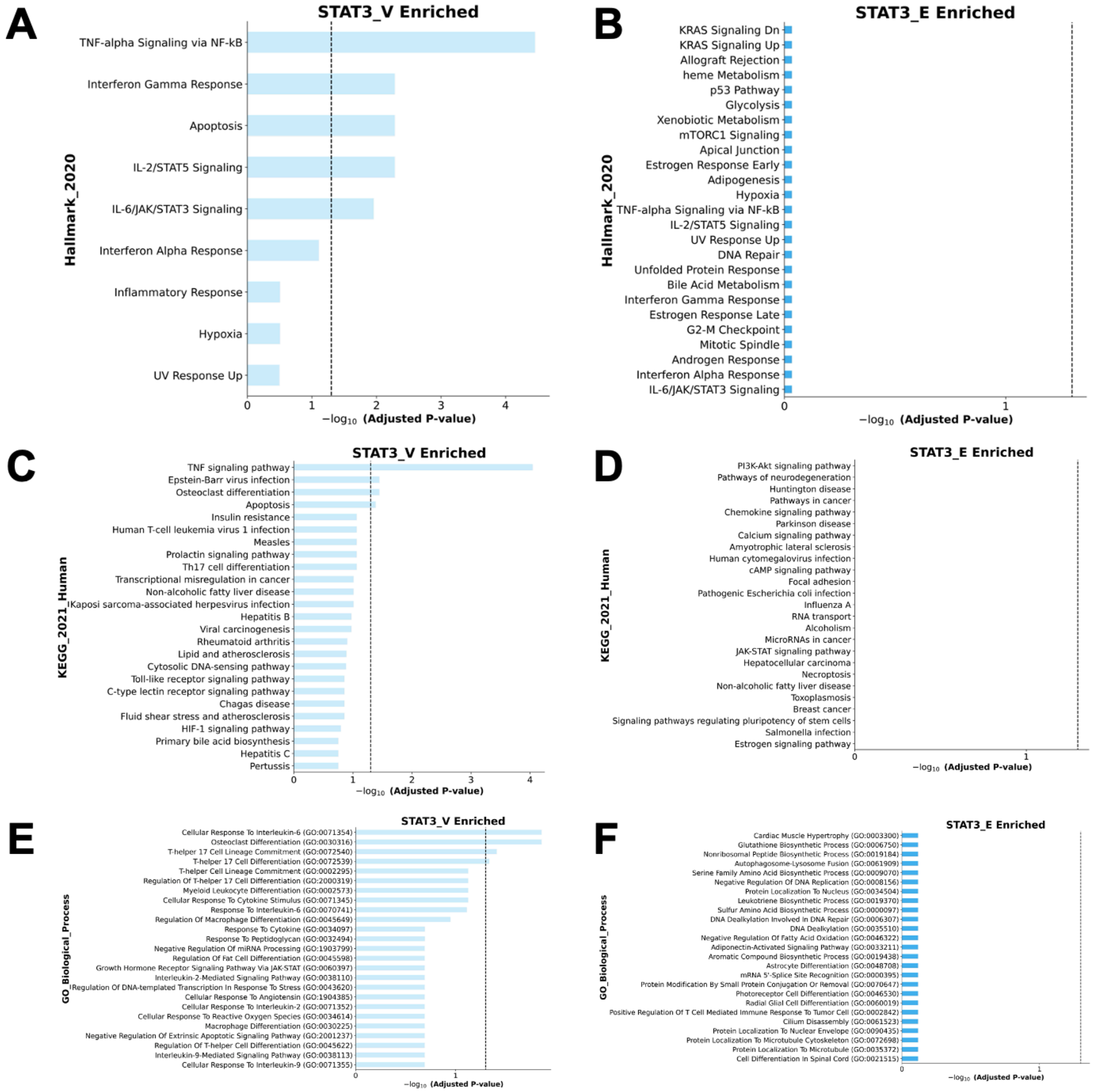


G

#### Vehicle

| Gene_set | Term | Adj. P-value | Genes |
|---|---|---|---|
| KEGG_2019_Mouse | TNF signaling pathway | 7.13E-05 | SOCS3, IL6, CASP8, CSF1, IRF1, RIPK1, FOS, JUNB, RELA, ICAM1, CREB5 |
| KEGG_2019_Mouse | Epstein-Barr virus infection | 1.57E-02 | IL6, CASP8, GADD45B, H2-Q6, STAT3, RIPK1, CD247, JAK3, RELA, IRF9, ICAM1 |
| KEGG_2019_Mouse | Osteoclast differentiation | 1.57E-02 | CSF1R, SOCS3, NCF1, CSF1, FOS, JUNB, IRF9, RELA |
| KEGG_2019_Mouse | Apoptosis | 2.49E-02 | PARP3, CASP8, GADD45B, LMNA, RIPK1, FOS, RELA, MCL1 |

#### Entinostat

| Gene_set | Term | Adj. P-value | Genes |
|---|---|---|---|
| KEGG_2019_Mouse | TNF signaling pathway | 9.71E-01 | FOS, MAP3K14 |
| KEGG_2019_Mouse | Epstein-Barr virus infection | 9.71E-01 | STAT3, H2-OB, CD3D, MAP3K14 |
| KEGG_2019_Mouse | Osteoclast differentiation | 9.71E-01 | FCGR1, FOS, MAP3K14 |
| KEGG_2019_Mouse | Apoptosis | 9.71E-01 | LMNA, FOS, MAP3K14 |

**Figure S6: STAT3 ChIP-seq GSEA pathways enrichment of treatment specific overall peaks**. **(A)** Hallmark_2020 GSEA enrichment barplot by enrichr method showing STAT3-vehicle enriched and **(B)** STAT3-Entinostat enriched pathways from STAT3 ChIP-seq of J774M cells. **(C)** KEGG_2021_Human GSEA enrichment barplot by enrichr method showing STAT3-vehicle enriched and **(D)** STAT3-Entinostat enriched pathways from STAT3 ChIP-seq of J774M cells. **(E)** GO_Biological_Process_2025 GSEA enrichment barplot by enrichr method showing STAT3-vehicle enriched and **(F)** STAT3-Entinostat enriched pathways from STAT3 ChIP-seq of J774M cells. **(G)** Tables of genes in differentially STAT3-enriched pathways from KEGG_2019_Mouse pathways analyses. Dotted line = p-adj. value cutoff of 0.05.

**Supplemental Figure 7**

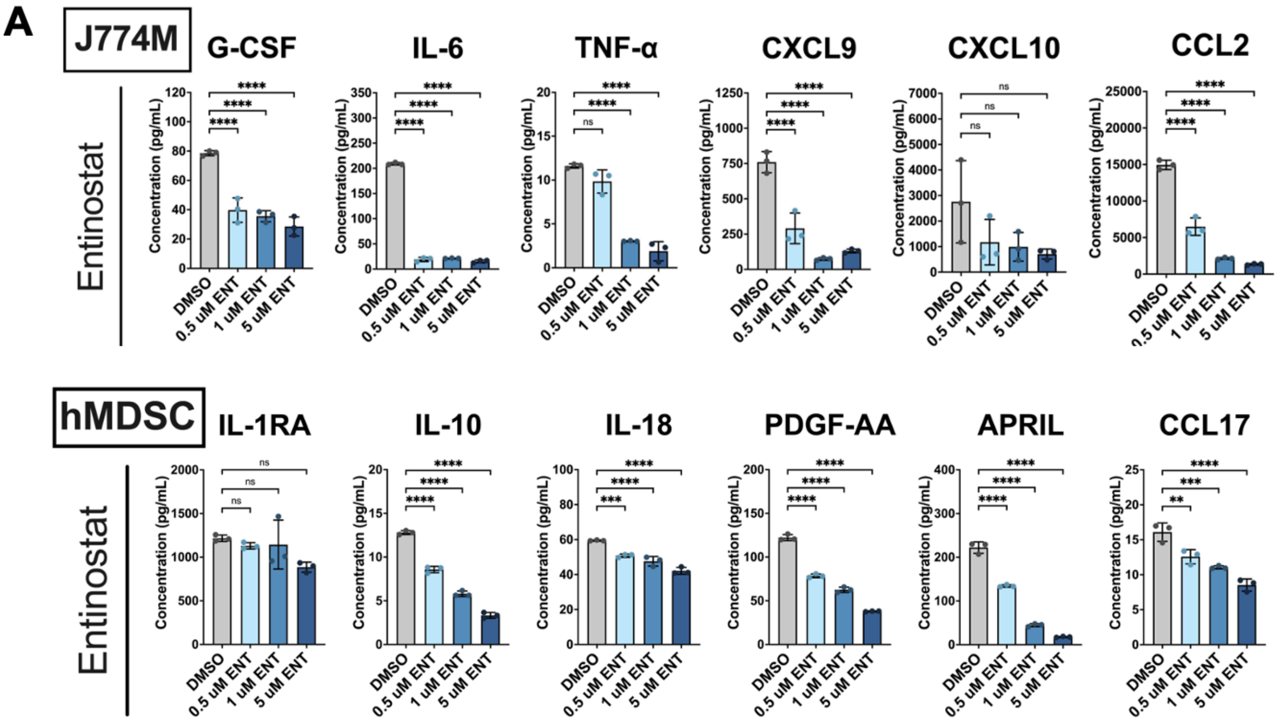


**Figure S7: Class I HDAC inhibition decreases suppressive cytokine secretion**. **(A)** ELISAs of supernatants from J774M (top, n=3) and hMDSCs (bottom, n=3) collected after 72 hours of Entinostat treatment. One-way ANOVA, ns = not significant, ** $p<0.01$, *** $p<0.001$, **** $p<0.0001$.
